# Desiccation plasticity and diapause in the Argentinian pearlfish *Austrolebias bellottii*

**DOI:** 10.1101/177386

**Authors:** Tom J M Van Dooren, Irma Varela-Lasheras

## Abstract

**Background:** The annual life history strategy with diapauses evolved repeatedly in killifish. To understand their and to characterize their variation between species, patterns of desiccation plasticity seem central. Plasticity might have played a role in the origin of these developmental arrests, when annual fish evolved from non-annual ones. The consequences of desiccation on survival and developmental rates of embryos of annual fish are poorly known. Using detailed demographic modelling of embryonal life histories, we investigate plasticity for desiccation in the Argentinian pearlfish *Austrolebias bellottii*. The treatment protocol contains changing environmental conditions with successive phases of mild desiccation and rewetting.

**Results:** We observed no clear diapause II and thus no increased incidence caused by mild and prolonged desiccation. Embryos arrest development in the pre-hatching stage (DIII) or in the dispersed cell phase (DI) irrespective of environmental conditions. There are limited effects of desiccation on survival, limited developmental delays and an acceleration of development into the pre-hatching stage. We found significant parental variance components on developmental rates, but hardly any effect of parental age. Hatching probabilities increased with age, when embryos had been in air at 100% RH and increased further when embryos were rewetted a second time after a two month interval.

**Conclusions:** Mild desiccation and rewetting affect survival, rates of development and hatching probability, but not the fractions of embryos that arrest development in particular stages. We can conclude that the incidences of diapause have become relatively independent of the occurrence of mild desiccation, as if they have become assimilated. In contrast to the responses to mild desiccation observed in the non-annual rivulids, *Austrolebias* accelerates development into the pre-hatching stage.

## Introduction

Diapause, where the progress of development is interrupted or halted, is a trait used to explain that evolution can favour delaying strategies and bet-hedging (Philippi and Seger 1989, Stearns 2000, Evans and Dennehy 2005). Diapause is not an instantaneous, easily reversible response to adversity (Danks 1987), but a developmental and metabolic arrest that can also occur in the absence of adverse environmental conditions. A decrease in metabolic activity during diapause allows embryos to survive such periods for long. These two main characteristics differentiate diapause from other types of developmental arrest or delaying strategies such as quiescence (which only happens in the presence of adverse environmental conditions) or delayed hatching (which does not encompass a metabolic arrest and is eventually deleterious for the embryo, Darken et al. 1998). Individual variability in such strategies can have components of bet-hedging and (transgenerational) plasticity together with genetic variation (Haccou and Iwasa 1995, Van Dooren and Brendonck 1998, Furness et al. 2015a, Polacik et al. 2017), adapting trait determination and rates of development to regimes of environmental variability. The study of individual life histories where diapause can occur therefore presents an excellent opportunity to link proximate drivers and properties of developmental processes with demography and long-term fitness or even phylogenetic evolutionary processes of species selection (Helmstetter et al. 2016).

Annual killifish are oviparous fish species from the suborder Aplocheiloidei (Berois et al. 2015) which use diapause to persist in temporary ponds of Africa and South America. These ponds dry out seasonally, killing all adults and juveniles present. Diapausing embryos buried in the soil survive the dry period and can hatch when the opportunity presents itself. This “annualism” life history has evolved repeatedly (Furness at al. 2015b, Helmstetter et al. 2016). Because fish eggs are transparent, the developmental dynamics of these embryonal life histories can be studied well in a non-invasive manner. Diapause in annual killifish can occur in three developmental stages (Wourms 1972a,c). Diapause I occurs between epiboly and the onset of embryogenesis, diapause II in the long-somite embryo, and diapause III occurs when the embryo is completely developed and ready to hatch. Diapause I takes place in a unique developmental stage after epiboly where the blastomeres individually disperse, and diapause II can coincide with a delay in the development of anterior structures with respect to the rest of the body (Podrabsky et al. 2010, Furness et al. 2015b). It is currently impossible to synonymize annualism with the occurrence of any of the separate three diapauses, because data on their occurrence and obligate or facultative character are lacking for many annual fish species. Among annual species studied thusfar, the presence of diapause I, II, and III and the different environmental cues that affect their onset, duration, and termination seem variable (Wourms 1972a,b,c, Markofsky and Matias 1977, Markofsky et al. 1979, Inglima et al. 1981, Matias and Markofsky 1984, Levels and Denuce 1988, Furness et al. 2015b).

Wourms (1972c) and Varela-Lasheras and Van Dooren (2014) have suggested that the evolution of plasticity has played a role in the evolution of the diapauses in annual fish and therefore that they might have originated in a scenario of genetic assimilation (Waddington 1942). Such a scenario is characterized by a temporary increase in the strength of phenotypic plasticity (reaction norm slope, Lande 2009), which then becomes reduced again or disappears. We previously found that non-annual Rivulids slow down early development in response to mild desiccation, and that hatching can be delayed over a prolonged period (Varela-Lasheras and Van Dooren, 2014; see also Furness 2015). Embryos with delayed hatching and decreased activity responded to a cue which provokes hatching in annual killifish such that the hatching delay observed showed several characteristics of diapause III. According Furness (2015), true annualism involves the occurrence of diapause I and II, and delayed hatching and some desiccation resistance would be commonly present in annual and non-annual species that encounter edge or marginal habitats with a risk of desiccation. To understand the evolution of the diapauses in killifish and to characterize variation between species, patterns of desiccation plasticity therefore seem central.

For annual killifish, however, there are no data allowing a detailed analysis of rates of development and survival in different developmental stages during and after a desiccation period. Podrabsky et al. (2001) found that embryos of the annual *Austrofundulus limnaeus* in diapause II can survive desiccation when air relative humidity (RH) is below 85% for over a month, whereas embryos in other stages did not survive for more than two weeks. Mortality was low and differences between developmental stages seemed small to negligible when RH was 85% or higher. This study did not assess effects of desiccation on the further development of the embryos that experienced desiccation and embryos were not returned to more humid conditions. Wourms (1972c) contains a brief statement that eggs of the Argentinian pearlfish *Austrolebias bellottii* (Steindachner 1881), an annual killifish from the sister taxon of non-annual rivulids, enter diapause I and II more often when subjected to partial desiccation, but data were not presented.

To investigate responses to desiccation of an annual rivulid related to the non-annuals investigated previously, we exposed embryos of the Argentinian pearl killifish *Austrolebias bellottii* in different stages of development to a period of mild desiccation and analyzed the effects on stage-specific survival and developmental rates, and on diapause incidence. Embryos were rewetted after desiccation, to investigate effects on hatching probability, and lagged effects on the further life histories of unhatched embryos after rewetting. The approach used to analyse the data is akin to the modelling proposed by Atchley et al. (1994), in that we mainly investigate and model rates of development (Varela-Lasheras & Van Dooren 2014). However, this is combined with modelling of survival patterns to obtain an integrative demographic model for the developing individuals.

## Methods

### Housing and data collection

The twenty-eight individuals of *Austrolebias bellottii* used in this experiment (14 males and 14 females from different family lines) were descendants from a population of 50 individuals collected in Ingeniero Maschwitz, Argentina in 2004 and hatched from eggs collected in 2008. There were two groups of pairs with an age difference of about 4 months (hatched two or six months before the start of egg collection, on April 20, 2010). Pairs were kept in 20-L aquaria in a climate room at 19°C with a 12-hour light/dark cycle. The aquaria contained some plants (e.g. *Elodea densa*, *Vesicularia dubyana*, *Lemna minor*) and a black, plastic 2L container half-filled with boiled coco peat where the fish could hide and lay eggs.

The day before collecting eggs, a clean container with 300 ml glass beads and 300 ml peat granules (both sieved and boiled) replaced the other container (as in Moshgani and Van Dooren 2011). Twenty-four hours later, this container was removed, the eggs collected and placed individually in wells of a 24-well plate with 1 mL of UV-sterilized tap water (DUNEA Leiden, The Netherlands) per well. Each clutch was divided over two different multiwell plates and placed in a different climate room with a temperature difference of 1 degree between them (19.5C vs. 20.5C). Next to the control treatment, which contained eggs continually incubated in water, we exposed the eggs to different “desiccation” treatments, limited to the range where expected survival of most stages would allow developmental responses to be observed (Podrabsky et al. 2001). We set up desiccators with demineralized water and saturated salt solutions of KNO_3_, NH_4_H_2_PO_4_ and KCl that would generate air humidities from 100 % relative humidity (Water), 94-93%, 93% and 86 %, respectively (O’Brien 1948). The humidity levels were verified with dataloggers (Gemini Tinytag PlusII, https://www.geminidataloggers.com) which could only confirm for KCl that the relative humidity was below 100% and at 85%. Some loggers might have failed to capture near 100% humidities reliably, due to wear on the probe or limited precision at high humidities, or some solutions were insufficiently saturated and did not produce the expected regime. For this reason, we decided to analyse differences between the different desiccation regimes with categorical variables and not by means of regressions on RH levels. Developmental states were determined minimally once per week for each individual embryo, except for one longer interval in summer (Figure 1). Eggs were transferred to the different desiccation treatments at varying ages between 17 days and 50 days. The desiccation treatments lasted between 51 and 94 days. Treatments and control were terminated by replacing the water in the wells with a mix of tap water, demineralized water and peat extract to promote hatching (Varela-Lasheras and Van Dooren 2014), and this event was followed-up over successive days. Alevins that hatched were removed from their well on the same day. Approximately thirty days later the water in the wells was replaced with sterilized water again for the eggs that had not hatched and there were no further observations made during a period of 61 days. The hatching procedure was then repeated (112 to 119 days after the first rewetting) and hatching scored again. In all, embryos experienced environmental sequences with up to four changes and were followed for up to 244 days.

**Figure one.**
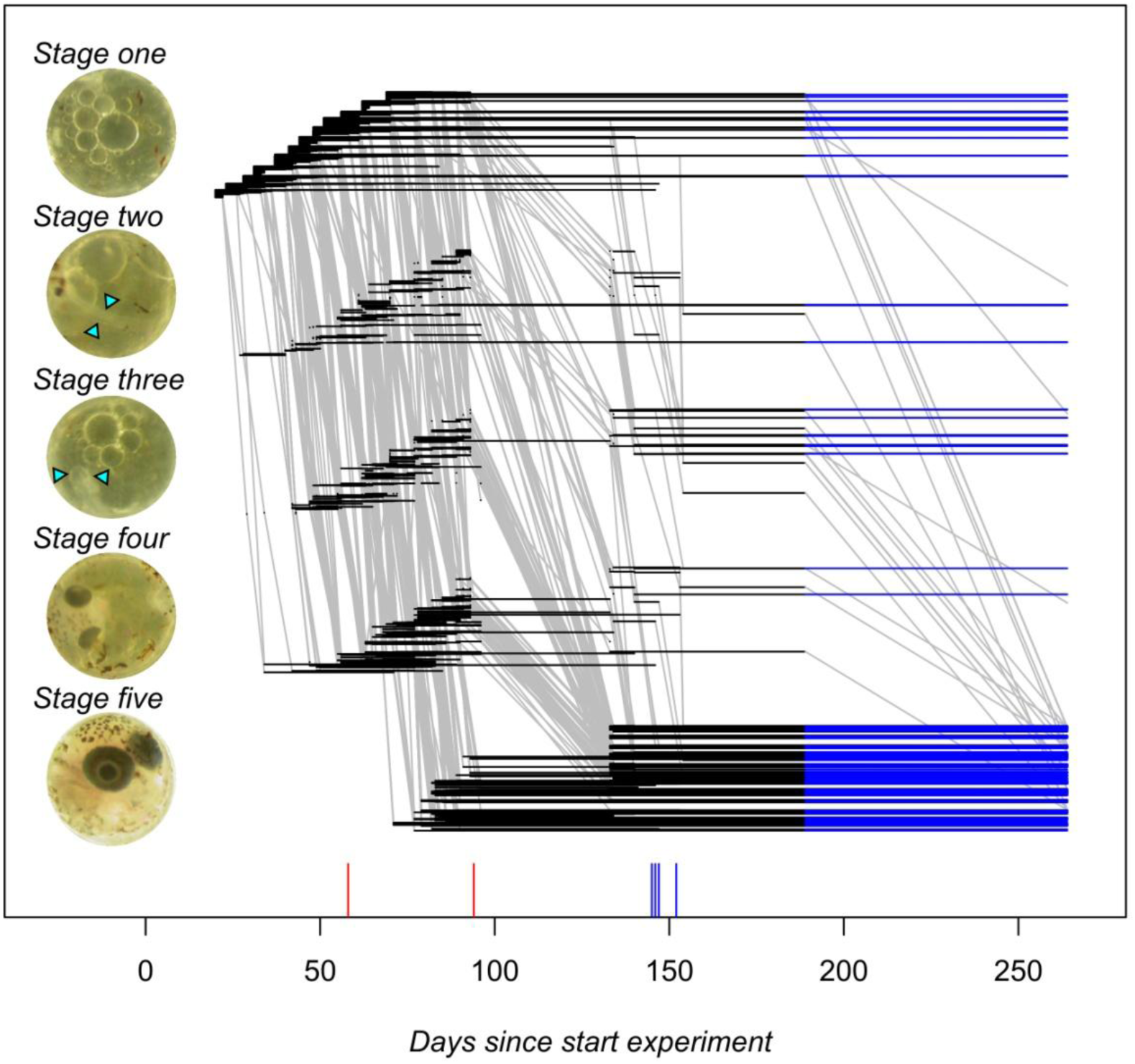
An overview of the “raw” data. Embryos were classified into five developmental stages as explained in the text, or as dead (this absorbing state is not represented). Per egg and stage, a horizontal line is drawn between the days of the first and last observation in that stage. Eggs are ordered within stage according their collection day. A blue line is drawn for eggs that were observed in the same stage at second rewetting (on day 264) and at the end of the period where regular observations were made (day 189). Developmental transitions are drawn as grey lines. Tthey connect the last observation in a stage with the first observation in the next stage where the egg was observed. Note that no observations were made between days 96 and 133. In the insets depicting the different developmental stages, small triangles indicate the location of the embryo when not clearly visible.

Based on previous studies (Wourms 1972a, Varela-Lasheras and Van Dooren 2014), individual state was scored as “dead” or as being alive in one of five developmental stages that can be distinguished well (Fig. 1). Briefly, embryos were assigned to stage one when collected. A neural keel and the first somites appearing delineate the start of stage two, the presence of optic cups the transition to stage three and the pigmentation of the eyes the transition to stage four. Finally, stage five starts when the embryo completely surrounds the yolk sac and ends when the fry hatches or dies. For embryos in stages four and five, we noted at each observation whether the tail coiled over the left or right side of the head.

### Statistical analysis

Developmental life histories consist of time-to-event data. The age intervals embryos spend in each developmental stage and environment before either moving into the next stage, hatching, or dying are the durations on which our analysis was based (Fig. 1). Events (deaths, developmental transitions) are assumed to have occurred just before they are observed. The data were analyzed using Cox proportional hazard models with random effects, using libraries survival and coxme in R (Therneau and Grambsch 2000, Therneau 2015). We modeled all transition rates per stage separately, that is, the mortality rates per stage and per-stage developmental transition rates to the next stage (Varela-Lasheras & Van Dooren 2014). The models per stage in the main text combine all durations in the different environmental conditions, so that effects of environmental conditions can be tested by removing them from the model. In the supplement, results are presented for models where the durations were analysed separately per environment (Control/Desiccation Treatments/Rewetted) or censored before the long inter-observation summer interval. In some environment/stage combinations, we had very few individuals at risk and therefore either did not fit any separate models to the data, or Cox models with fixed effects only. We also analysed some zero-one responses or proportions of states present at specific time points using logistic regressions. In order to give an overview of individuals in different stages depending on age, we used multi-state estimation of fractions per stage and made plots of these fractions using the mstate library for R (de Wreede et al. 2011).

**Survival.** Per stage, we first fitted a Cox model with period (control “Wet”/ desiccation “Dry” /”Rewetted”) and salt treatment (“H2O” / “KNO3” / “NH4H2PO4” / “KCL”) fixed effects, their interactions and temperature and parental age (categorical, “Young” / “Old”) fixed effects. For stages two to five, we added the duration spent in the preceding stage as a covariate. The model contained a parental pair random effect, a random effect of collection date and a plate effect. This model was then simplified by removing effects that were not significant in a chi-squared likelihood ratio test. We started model simplification by inspecting the significance of random effects, followed by tests on the fixed effects. We sequentially tested and deleted non-significant effects in a fixed order: first the tests for pair variation, collection date, plate variation; then previous duration effects, salt treatments, the period effect, temperature and age effects. When an effect was significant in a likelihood ratio test but the parameter estimate had a very large standard error or there were data lacking to estimate a subset of parameters, we also removed that effect from the model or analysed a subset of the data. We encountered numerical problems fitting models with random effects that had different variances per period. Therefore we could not test directly whether these variances depend on the environmental state and analysed the data per period separately as well. The models in the Supplement fitted to the data per separate stage and period (i.e. without period effects) underwent the same model selection procedure.

**Development.** The data on developmental rates were analyed starting from models containing the same maximal models as for survival and we tested effects in the same order. For each separate analysis per developmental stage, age within stage is used as the appropriate time scale. Spontaneous hatching from stage five was modeled as a developmental transition, while hatching induced by rewetting caused censoring.

**Fraction in diapause.** We investigated whether the distribution of embryos across stages at the moment of rewetting and at the end of the experiment depended on previous individual history. We fitted generalized linear models (glm’s) to the data, analysing the multinomial data distribution with a set of binary logit models, each for the fraction of embryos in an earlier developmental stage relative to the number in stage five (baseline category logit models, Agresti 2013). Mixed binary models did not converge well here, we therefore restricted the analysis to fixed effects of desiccation treatment, time spent in the treatment and their interaction, and added age and temperature main effects. We used the inverse of the time spent in a treatment as an explanatory variable such that the intercepts per treatment became the expectation for when embryos would have been in the treatment for an infinite amount of time. Model selection was carried out using likelihood ratio tests, as the data were 0/1 individual responses with no need for overdispersion corrections.

**Coiling.** For embryos in stage four and five we repeatedly observed the direction in which they coiled their tail over the head, and used the rate of change in direction as a proxy for activity (Varela-Lasheras and Van Dooren 2014). We estimated how the probability of a transition between left and right-coiling depended on treatment, parental pair, and time spent in stages four or five. We used glm’s for the occurrence of a change in coiling (positive response) and with a complementary log-log link, so that by means of an offset we could correct for the duration between observations (Harney et al. 2013).

**Hatching.** The probabilities of hatching when water with peat extract was added are reported. We analysed these probabilities of hatching using binary logit generalized linear models. Individuals that hatched within three days of adding peat water were scored as a positive response, the ones that remained in stage five as non-hatchers. Individuals that were found dead in this time interval were not included in the analysis, as it was difficult to determine whether they had actually hatched or not. This model contained as explanatory variables effects of temperature, collection date, date where hatching water was added, desiccation treatment, parental pair, total age, age in stage five, time spent in treatment and interactions of desiccation treatment effects with all three time variables. Model selection was carried out using likelihood ratio tests.

## Results

We collected and incubated 2161 eggs in total, 539 embryos were subjected to desiccation treatments (including air at 100% humidity) and 599 embryos from the Control and desiccation treatments were rewetted. The median number of observations per embryo was 16 and the maximum 31. Table 1 shows that the desiccator with KCl contained fewer embryos at risk than the other treatments. We decided to do that to minimize individual losses at the strongest level of desiccation in this experiment.

**Table 1.**
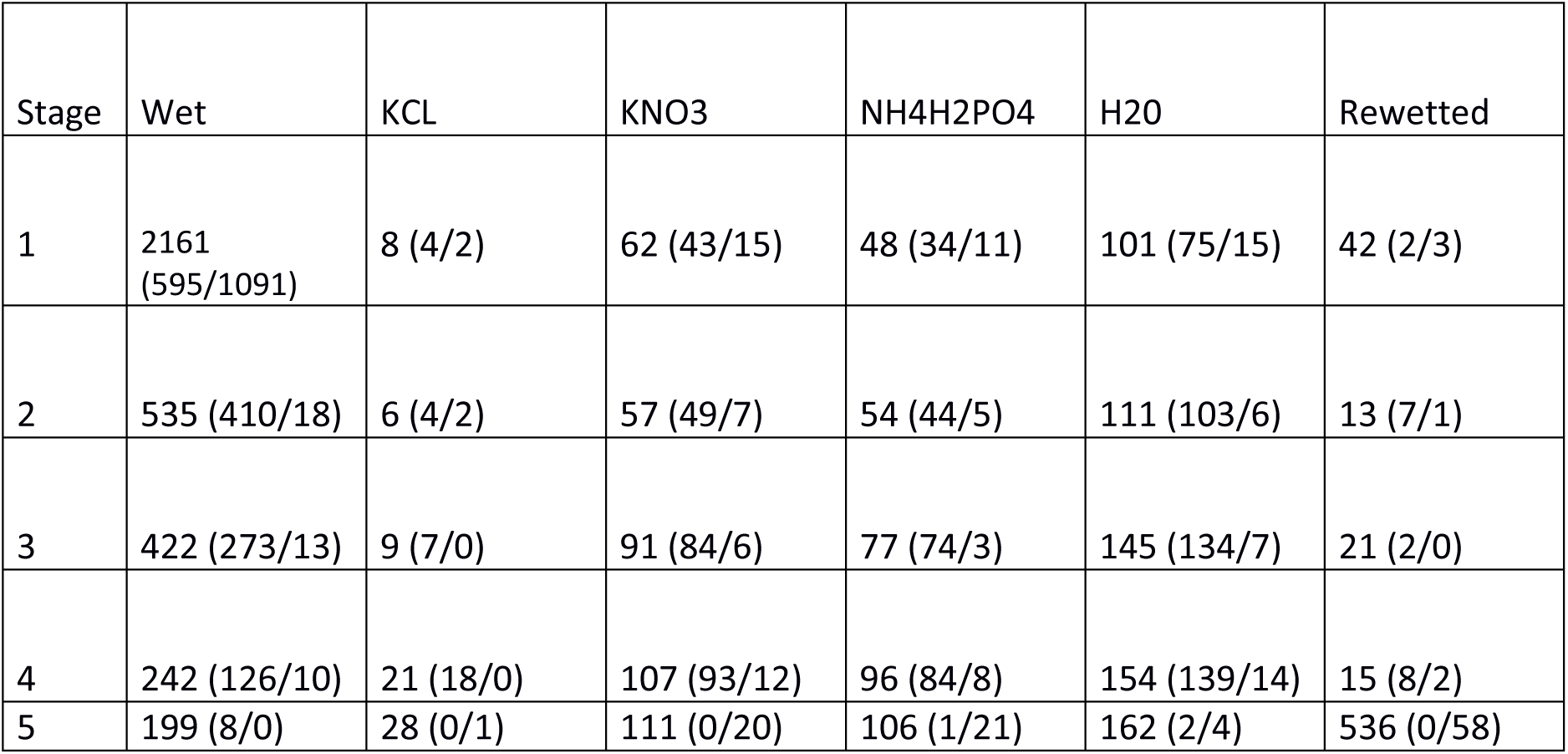
Number of individuals at risk (number of developmental transition events, number of deaths within stage) per stage and period/salt used for desiccation.

| Stage | Wet | KCL | KNO3 | NH4H2PO4 | H2O | Rewetted |
| --- | --- | --- | --- | --- | --- | --- |
| 1 | 2161<br>(595/1091) | 8 (4/2) | 62 (43/15) | 48 (34/11) | 101 (75/15) | 42 (2/3) |
| 2 | 535 (410/18) | 6 (4/2) | 57 (49/7) | 54 (44/5) | 111 (103/6) | 13 (7/1) |
| 3 | 422 (273/13) | 9 (7/0) | 91 (84/6) | 77 (74/3) | 145 (134/7) | 21 (2/0) |
| 4 | 242 (126/10) | 21 (18/0) | 107 (93/12) | 96 (84/8) | 154 (139/14) | 15 (8/2) |
| 5 | 199 (8/0) | 28 (0/1) | 111 (0/20) | 106 (1/21) | 162 (2/4) | 536 (0/58) |

**Survival.** The data represented in Figure one are first used to draw survivorship curves (Figure two, A-E). For age intervals where these curves are steep, mortality rates are high. Where they become flat, mortality becomes neglible. Figure two shows that, overall, survival is highest for pre-hatching embryos (Fig. 2E) and lowest for the earliest developmental stage (Fig. 2A). What is also apparent is that the death rate decreases with age to become zero in most stages and treatment groups. By the age where all surviving embryos are in the rewetting phase, there are hardly any new deaths occurring. Table 2 summarizes which variables were retained in the models for survival per stage, For comparison, Table S1 does that for a dataset restricted to the observations made before the start of the single long interval in between observations (days 96-133, Fig.1). For stage two and stage five, there are significant fixed effects. Other treatment effects suggested by Fig. 2 were not significant. Embryos that are desiccated in stage two have an increased death rate (c.i. [0.11, 1.49]) and embryos which have previously spent a longer duration in stage one have a decreased death rate (c.i. slope [−0.12, −0.02]). In stage five, there is negligible mortality in the wet phase (Fig. 2 and Figure Surv5 in the Supplement). Death rates in the pre-hatching stage are increased by any of the desiccation regimes, even when eggs were in air with 100% RH (increase in mortality rate for H_2_O c.i. [0.33, 3.00])). Confidence intervals for mortality rates of KNO3 (c.i. [2.45, 4.81]) and NH4H2PO4 ([2.51, 4.87]) desiccation regimes overlap with H2O, for KCL it does not (c.i. [3.68, 6.31]). It needs to be noted that the order of mortality rates is according the intended strengths of desiccation. In the pre-hatching stage, mortality increases with temperature (c.i. [0.89, 1.81]). The supplementary material contains Table S3 and figures of survivorship curves per stage and per period (Wet/Dry/Rewetted). These results suggest that the lower survival in the driest desiccator was caused by a briefly elevated mortality at the start of the rewetting phase. There are no significant effects of parental age (Table 2). However, in stage five and when each period is analysed separately (Table S3), we recover parental age effects not found when the data are jointly analysed across periods. Pre-hatching embryos with older parents have lower mortality in the dry period (parameter estimate −1.49 (0.42)) but a higher mortality in the following rewetted period (1.43 (0.75)) resulting in no net overall effect.

**Table 2.** Survival analysis per stage across periods. “NS” indicates non-significant effects. Period × treatment interactions were never significant and therefore not listed. Significant effects are indicated by Chi-squared statistic and p-value. For the salts used, those which had significantly different parameter estimates from desiccators with water are given differently shaded cells.

| Stage | Random effects |  |  | Fixed effects |  |  |  |  |  |  |  |
| --- | --- | --- | --- | --- | --- | --- | --- | --- | --- | --- | --- |
|  | Parent | Date | Plate | KCL | KNO3 | NH4H2PO4 | Salts | Periods | Temp | Age | Preceding |
| 1 | 134.25<br>< 0.001 | 71.70<br>< 0.001 | 12.41<br>< 0.001 |  |  |  | NS | NS | NS | NS |  |
| 2 | NS | NS | NS |  |  |  | NS | 5.19<br>0.023 | NS | NS | 11.67<br><0.001 |
| 3 | NS | 9.08<br>0.003 | NS |  |  |  | NS | NS | NS | NS | NS |
| 4 | NS | 4.78<br>0.029 | NS |  |  |  | NS | NS | NS | NS | NS |
| 5 | NS | NS | NS | + |  |  | 157.02<br><0.001 | NS | 37.25<br><0.001 | NS | NS |

**Figure two.**
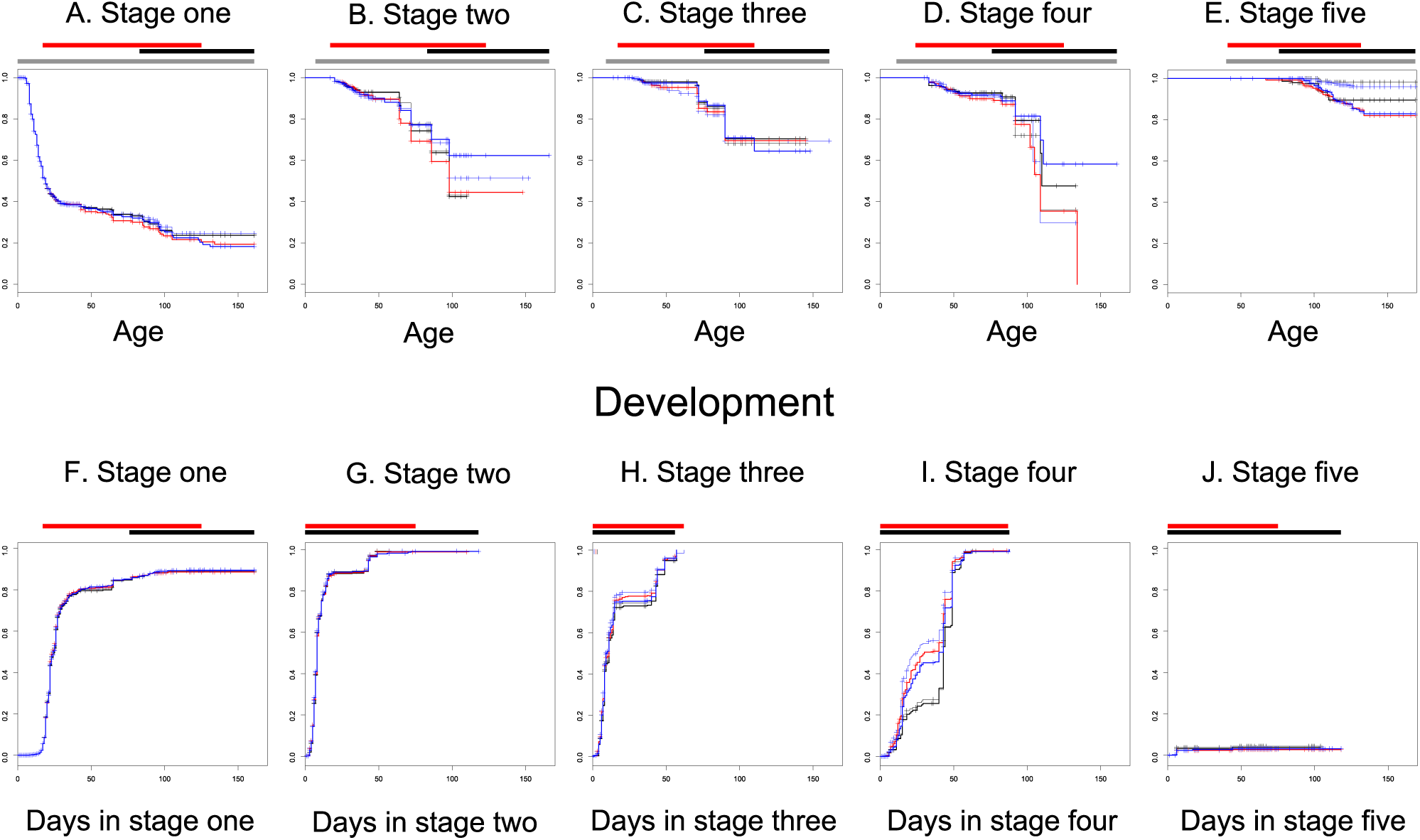
Survival and development represented with survivorship curves. Each panel contains the data for a specific developmental stage and has a separate survivorship curve per treatment group. At many instances, embryos within the same stage were distributed over different environmental regimes (control / dry / rewetted). Bars above panels indicate ages were embryos were in that stage (grey, survival), in the desiccation regime (red) or rewetted (black).

When we inspect the results on the random effects, then variances between parental pairs and between plates are significant in stage one only, while random effects of collection date are present in three out of five stages (stages one, three and four).

**Development.** Figure 2 F-J represents the proportions of embryos that have made a developmental transition depending on the time already spent in each stage. For cohorts of embryos in stages one and five, the rate of development or the probability of spontaneous hatching become zero before all individuals have made a developmental transition (Fig. 2F, J), showing that diapause I and III occur irrespective of mild desiccation. There is a small fraction of abortive hatchings from stage five. Diapause II is not observed as a levelling off in the fraction that made a transition (Fig. 2H), even with prolonged but mild desiccation. What we do see is that the curves for development in stage two, three and four all show a period of about twenty days between days 20 and 50 with fewer developmental transitions. We found out that this is due to the occurrence of the single long time interval between observations. When data were censored before that interval, there is no such period and we conclude that there are no slow/fast alternative developmental trajectories in these stages. For the shortened dataset and relative to the Control, embryos in saturated air develop significantly slower (-0.37, s.e. 0.17) from stage one into two. However, the effect is not clearly visible or strong in graphs, there are no significant effects for the other treatment levels and the effect is not detected when the complete data are analysed. We therefore denote it as spurious. Desiccation changed the developmental rate from stage two into stage three relative to wet conditions (Table 3 and parameter estimates H2O 0.27 (s.e. 0.23); KNO3 −0.21 (0.25); NH4H2PO4 −0.57 (0.26)); KCL 1.51 (s.e. 1.06). This pattern of estimates suggests a non-linear effect of desiccation intensity, but that suggestion strongly depends on the estimate for KCL which has large sampling variance and is therefore the least reliable. The pairwise difference from the control group is only significant for the NH4H2PO4 treatment. These differences are not discernible in Fig.2G where individuals in Control, desiccation and rewetted regimes are pooled for most of the time points. However, they are visible in Figure SDev2 in the Supplement. Individuals that spent more time in stage one have a slower development rate from stage two into tree (Table 3, slope −0.030 (0.006)). Figure Dev4 in the supplement suggests that for developmental rates from stage four into five, KCL and the control groups have a lower rate of development than other treatments. The mixed Cox models confirm this, with development into stage four significantly increased relative to Control in a desiccator with H2O (1.73 (0.41)) KNO3 (1.49 (0.43)) and NH4H2PO4 (1.18 (0.46)). The rate of development in stage four tends to be accelerated for embryos that have spent more time in the preceding stage (slope 0.044 (0.024), Table 3, TableS2, S4). The rate of abortive and spontaneous hatchings from stage five is significantly smaller for embryos that are in the desiccators (parameter estimate −2.24 (0.72)) and there are no abortive hatchings in the rewetting phase (Figure SDev5 in the Supplement).

**Table 3.** Analysis of developmental rates per stage, across periods. “NS” indicates non-significant effects. Period × treatment interactions were never significant and therefore not listed. Significant effects are indicated by Chi-squared statistic and p-value. For the salts used, those which had significantly different parameter estimates from desiccators with water are given differently shaded cells. For stages written in bold, we did not include time intervals in the rewetted period in the analysis.

| Stage | Random effects |  |  | Fixed effects |  |  |  |  |  |  |  |
| --- | --- | --- | --- | --- | --- | --- | --- | --- | --- | --- | --- |
|  | Parent | Date | Plate | KCL | KNO3 | NH4H2PO4 | Salts | Periods | Temp | Age | Preceding |
| 1 | 79.68<br><0.001 | 8.9<br>7<br>0.003 | NS |  |  |  | NS | NS | NS | NS |  |
| 2 | NS | 8.55<br>0.003 | 4.53<br>0.033 |  |  |  | 10.34<br>0.016 | NS | NS | NS | 30.31<br><0.001 |
| 3 | NS | NS | 7.46<br>0.006 |  |  |  | NS | NS | NS | NS | NS |
| 4 | 6.64<br>0.010 | 7.34<br>0.007 | 4.61<br>0.032 |  |  |  | 9.56<br>0.023 | NS | NS | NS | 3.21<br>0.073 |
| 5 | NS | NS | NS |  |  |  | NS | 11.39<br>0.003 | NS | NS | NS |

We find significant variation between pairs for the first and fourth stage (Table 3). Plate effects for developmental rates occur in slightly more developmental stages than for survival rates (Table 3, S2, S4).

**Multistate overview** We made an overview figure of fractions of embryos in different states across time with different panels for the different periods (Figure 3). The panels again show the decreases in rates when embryos have been a while in a treatment – the boundaries between states become more horizontal. The amount of events is overall small in the rewetting period. Among the individuals that didn’t hatch, stage five embryos are the most numerous at the end of the experiment, suggesting that in the environments we investigated, diapausing individuals are mainly of diapause type III after some time. Note also that among the rewetted embryos, there is a small fraction of individuals in stage three. This could not be seen in Fig. 2H, where the age-within-stage intervals for desiccated and rewetted individuals overlapped nearly completely for that stage.

**Figure three.**
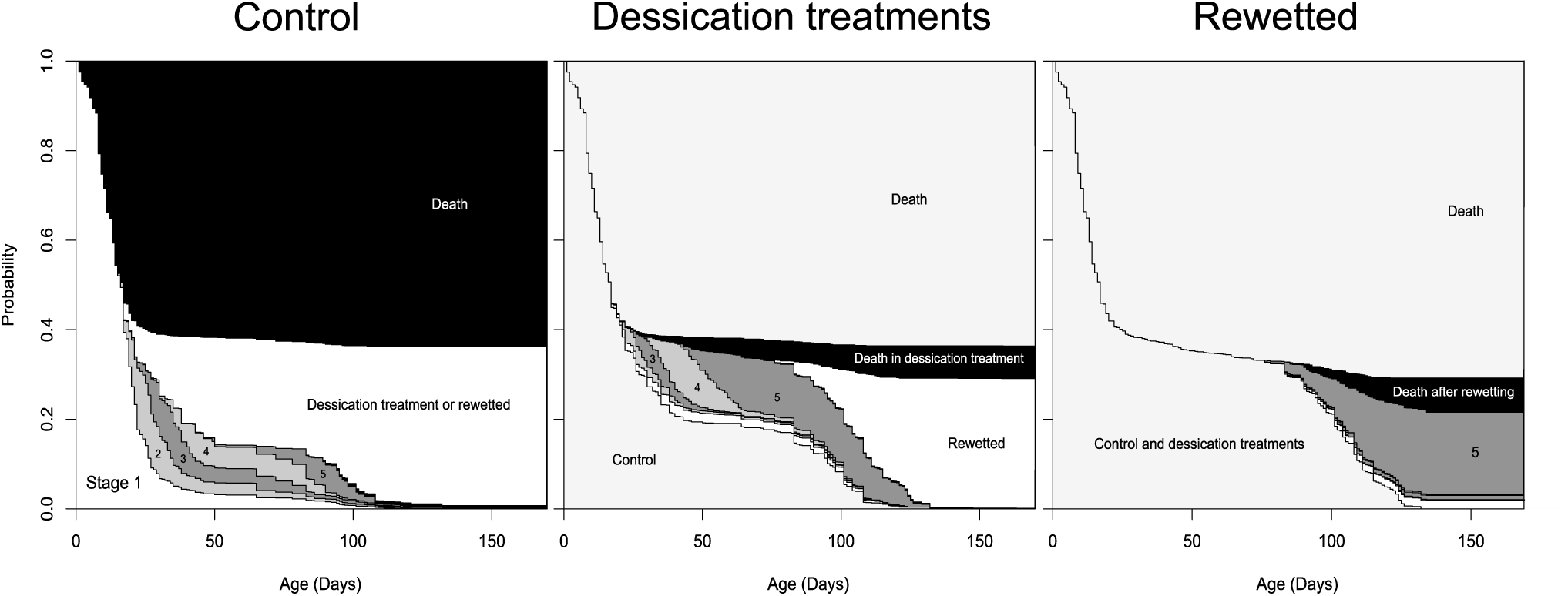
Proportions of embryos in different states (stages of development, death) in dependence on their age. Proportions were estimated using multistate survival analysis. Wet, dry (humid air) and rewetted periods are detailed in separate panels. Each panel only shows the fractions of embryos in the different developmental stages for the focal environmental regime. In the left panel, embryos that entered the dry and rewetted regimes are categorized separately as being in that state. In the middle panel, the distribution of embryos in the dry regime over different states (including death) and the fraction that entered the rewetted state are shown. In the right panel, the distribution of embryos over stages in the rewetted regime are detailed.

**Distributions across stages.** At first rewetting, 42 individuals were in stage one, 6 in stage two, 19 in stage three, 7 in stage four and 525 in stage five. We only analysed fractions in stages one and three relative to stage five using glm’s. For the fraction of embryos in stage one relative to stage five at rewetting, we found a trend for the *time in treatment* × *treatment* interaction (chisq 10.79, df = 5, *p* = 0.06). This suggests that the speed at which the proportion of embryos in stage one stabilizes relative to the fraction in the pre-hatching stage differs between treatments. However, there are no intercept terms that differ significantly. There is therefore is no parental age effect on the eventual fraction in stage one, if the time spent in treatment is taken into account. The fraction of embryos that are in stage three at rewetting is smaller for older parents (effect estimate −2.06 (s.e. 0.53); chisq 17.8, df = 1, *p* < 0.001). The parental age effect on the fraction in stage three persists when time in treatment is included in the model. For the distributions over stages at the end of the observation period, we find the same effects. We can interpret this at most as different speeds within treatments at which the remaining individuals settle in diapauses I and III, but no consistent treatment differences in the eventual fractions. When we represent the distribution over stages at first rewetting in a graph (Figure 4), it can be seen that the older parents contributed more individuals overall and in particular in the pre-hatching stage, as they produced larger clutch sizes sooner. At the end of the period where we observed individual state regularly, 137 of the stage five individuals had hatched or died.

**Figure four.**
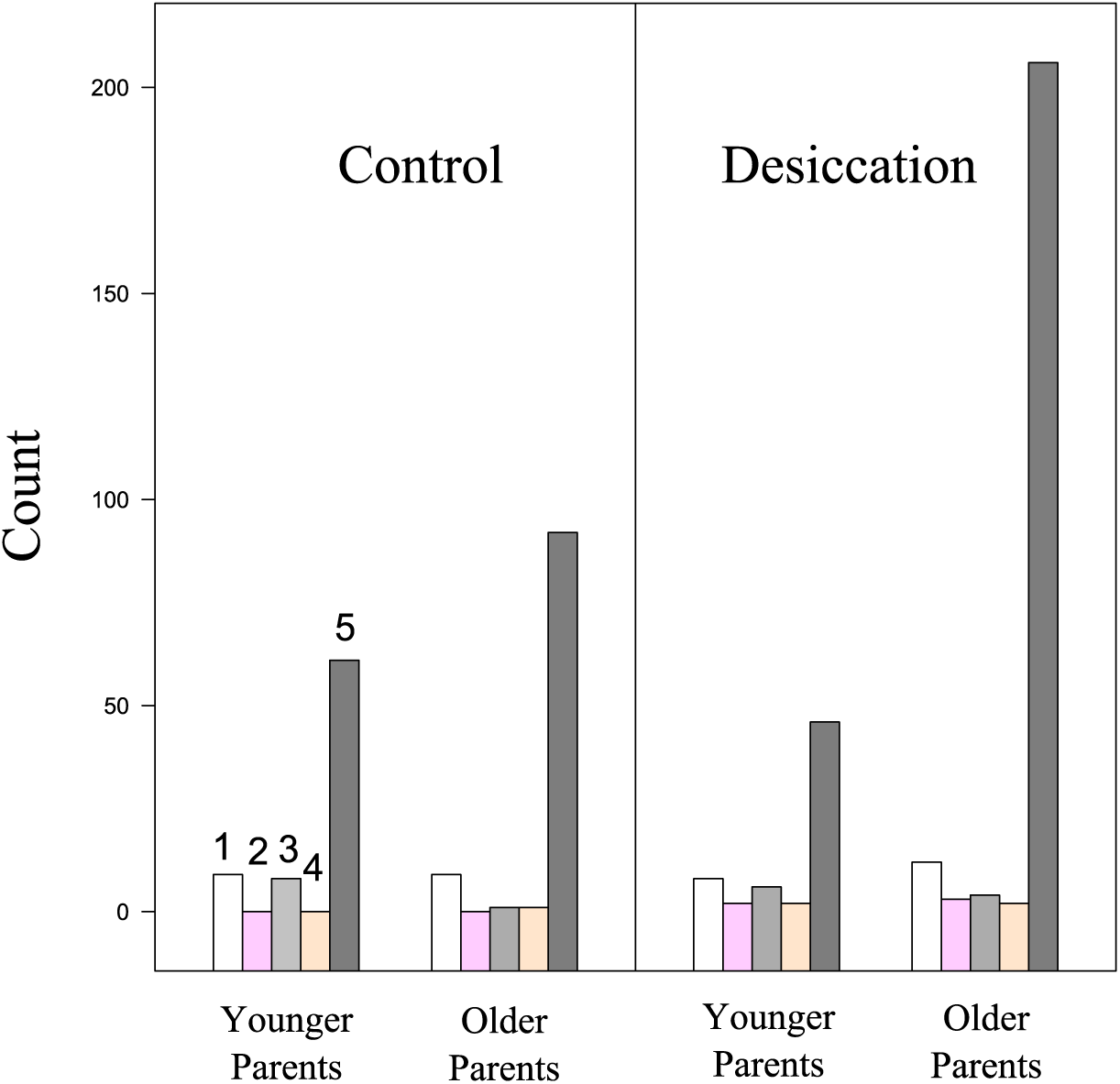
Distributions over stages at rewetting, for embryos in control (Left Panel) and desiccation regimes (Rigt Panel). Embryos of older and younger parental pairs are represented separately. There is a bar for each developmental stage we scored. These are ordered from one to five, from left to right.

**Hatching** On average, the hatching probability of the Control group where embryos were in water at all times is 12%. For the H2O group it is 27%, in theKNO3 desiccator the hatching probability is 13%, from NH4H2PO4 12 % and there was no hatching in the KCL dessicators. Mixed models for hatching probabilities did not converge well, and we refitted them as glm’s. For the earliest set of days where peat water was added, we found significant effects of collection date (chisq(11) = 40.29, *p* < 0.001), a time in treatment × treatment interaction (chisq(5) = 29.46, *p* < 0.001), and a negative parental age effect (chisq(1) = 5.08, *p* = 0.02). In the H2O treatment, the intercept term is increased relative to the control (H2O 6.33 (s.e. 3.06)).

When we inspected embryos before the second rewetting (Table 4), a fraction of the eggs had developed further starting from stages one to four. Other embryos had remained in the stages where they were two months before, which was the case for stages one, two, three and five. Survival did not differ significantly between stages. Overall survival was 83 % across this period. When peat water was added, 48% of the stage five individuals hatched. We found significant effects of temperature (chisq(1) = 5.02, *p* = 0.02) and of the date where the embryos were rewetted before (Chisq(3) =30.52 *p* < 0.0001). For embryos stored at 20.5 degrees, the hatching probability was reduced (parameter estimate effect −0.56 (s.e. 0.25)). Two first rewetting dates had significantly reduced hatching probabilities at the second rewetting (parameter differences approx. −1.5).

**Table 4.** Developmental stages at second rewetting. A two month-period separated the follow-up of embryos after a first rewetting from the second rewetting. Numbers of embryos are given for each combination of states at the end of the follow-up period and at the moment of rewetting.

|  | Stage at second rewetting |  |  |  |  |  |
| --- | --- | --- | --- | --- | --- | --- |
|  | 1 | 2 | 3 | 4 | 5 | Dead |
| State at end of follow-up after first rewetting |  |  |  |  |  |  |
| 1 | **21** | 1 | 1 | 0 | 5 | 11 |
| 2 somites | 0 | **3** | 0 | 0 | 1 | 1 |
| 3 head formed | 0 | 0 | **8** | 1 | 7 | 3 |
| 4 pigmented | 0 | 0 | 0 | **0** | 3 | 2 |
| 5 pre-hatching | 0 | 0 | 0 | 0 | **309** | 79 |

**Coiling.** We estimated rates of change of the tail coiling orientation using logistic regression, with a glm as in Harney et al. (2013). In stage four, we found a significant treatment × time in stage interaction (chisq(4) = 908.84, *p* < 0.0001) and differences between parental pairs (chisq(6) =13.59, *p* = 0.035), but none of the parameter estimates were significantly different from zero. When simplifying the model to a single effect of time within age, we found that it was non-significant.

In stage five, there is a significant treatment × age within stage interaction (Chisq(3) = 9.072, *p* = 0.0284), but again no separate parameters could be shown to be significantly different from zero.

For the group desiccated with NH4H2PO4 the probability tends to be lower (-1.48, 0.82, *p* = 0.07). Simplifying to a single covariate - age within stage five - we find that the probability of changing direction decreases with age in stage five (parameter estimate (-0.092 (s.e. 0.038)). According to this simple model, at age-witin-stage zero the probability to change coiling direction for stage five is 5% per day, twenty days into stage five it is 0.8% and at day forty 0.1 %. For stage four, the probability to change coiling direction is 7% per day.

## Discussion

We observe that irrespective of whether environmental switches were imposed, a general slowdown in rates of events and no further deaths nor developmental transition events were observed when the environment remained constant for a prolonged time. It is as if all rates become as good as zero and the developmental system slows down for an extended period of time, while mortality is arrested.

In the Argentinian pearlfish, embryos enter diapause I and III irrespective of the presence of mild desiccation and before transition rates become zero overall. Diapause II is at best rare and does not increase in frequency with desiccation. We do observe embryos that spend substantial periods in stages two and three, after water has been replaced two months before the second rewetting, but is unclear whether that is due to overall low rates of development in that period. Our data suggest that long observation periods combined with repeated environmental changes will be necessary in *Austrolebias* to determine whether diapause II truly occurs. Whether an embryo has been developing slowly through a developing stage can have effects on subsequent speeds of development, but the effects are not systematically magnifying the range of variation in timings. We observe an acceleration of development and growth from stage four into five with mild desiccation. For the pre-hatching stage, the rate at which coiling changes gradually decreased with age-within-stage, suggesting that overall levels of activity decrease during late development. This is in agreement with intensifying diapause III, while there there were no significant desiccation effects. We observed an increased probability of hatching when embryos have been in the 100% RH enviroment. When the hatching protocol is repeated two months later, there are no remaining lagged effects of the desiccation treatment and embryos hatch again with a higher probability. In our experiment that was at an age of about six months. Is desiccation not provoking any delayed effects? We found overall effects of desiccation when data were analysed across periods, so we cannot exclude that desiccation has delayed effects during rewetting. When data are analysed per period separately, desiccation effects are not demonstrable in the rewetting period, due to the low number of events observed there, as all processes have already slowed down.

We note that survival effects of desiccation were significant only for the pre-hatching stage, hence involving diapause III embryos.

### Annual vs. non-annual rivulids

Striking here in comparison with previous results on species from the non-annual rivulid sister group (Varela-Lasheras & Van Dooren 2014) is that passing through a desiccation period has limited effects and can accelerate development. In non-annuals, delays were early in development and desiccation provoked an increased incidence of delayed hatching (similar or equal to diapause III) in stage five in some species. In the annual, the speed of developing near head formation was slowed after rewetting for one desiccation level, or showing a positive response to desiccation late in development. Regarding early development in the first stage, we only found a significant effect in a reduced dataset and for the weakest desiccation level, casting doubt on its existence. The desiccation regimes in this study are much prolonged in comparison to Varela-Lasheras and Van Dooren (2014), where the median duration was eight days, with visible effects after one hour. In non-annuals, desiccation significantly increased mortality overall in embryos in stage three and four and decreased in *C. magdalenae* again after return to water. In the annual, developing embryos are quite resistant to the desiccation levels we applied as survival effects are limited to the pre-hatching stage and we could impose mild desiccation as a prolonged treatment. Note that there are no escape embryos in the sense of Polacik et al. (2017) that reach the pre-hatching stage within a month.

Both *Austrolebias* and non-annual rivulids show diapause III-like characteristics. This trait, very similar to delayed hatching, might be ancestral in the group. Slowing down during early development in non-annuals in response to desiccation could be mostly an immediate responses therefore quiescence, even while there are some lagged effects. If genetic assimilation of diapause has occurred in killifish via the evolution of plasticity to desiccation, then there has clearly been further plasticity evolution after the emergence of diapause-like characteristics. The plasticity now involves an accelerating effect.

The activity of pre-hatching annual embryos seems overall lower from the start in comparison to non-annuals where initially over 10% of individuals changed coiling per day. Also notable is that hatching probabilities were relatively low at the first rewetting and doubled at the second rewetting.

### The three diapauses

Diapause II might be rare in *Austrolebias*. According to Wourms (1972c) diapause I is common in South-American *Austrolebias* killifish, whereas diapause III is obligatory and diapause II facultative, as in Nothobranchius (Wourms 1972c, Furness et al. 2015a). This would render the first and last diapauses the most defining characteristics of the annual strategy for the *Austrolebias* genus. Several papers recently brought new data and ideas on the diapauses in annual fish. Furness (2015) believes diapause I to have a short duration in benign conditions and to be mainly brought about by harsh conditions. Diapause II, after the phylotypic stage, would contribute most to variability in developmental duration (Furness 2015). Podrabsky et al (2001) found that *Austrofundulus* embryos resisted desiccation best in this stage. Diapause II at low temperatures is not nearly obligate as in *Austrofundulus limnaeus* (Podrabsky et al 2010), but at best facultative in *Austrolebias*. Given the results here, it is certainly not the most prominent stage of arrest in annual fish as proposed by Furness (2015). In the Argentinian pearl killifish and in presumably benign conditions virtually all embryos showed behaviour classified as diapause III, a fraction diapause I, and diapause II is at best rare and potentially only occurs in older eggs.

Hatching of the stage five embryos was limited at the first wetting, the fraction hatching increased several months later. This calls into question the hypothesis that diapause III capitalizes on within-season hatching opportunities (Furness 2015), if the hatching response depends on age in this manner.

At first rewetting we had fractions of individuals in all stages. The stage three individuals might be in diapause II but flipped survivorship curves (Fig. 2H) suggest that there are no individuals remaining in stage two or three for long at that age. Our results are in agreement with Wourms (1973), that diapause II is not constitutive for this genus. Being more facultative than diapause III, it can not be equated to the annualism syndrome (Furness et al. (2015b). Diapause II shows strong plasticity in *Nothobranchius* (Furness et al. 2015a) and has negligible incidence in *Austrolebias* at a temperature where it is common in *Nothobranchius furzeri*. In Diapause II, development is arrested at a point where maintenance costs of the thusfar developed organs and structures might be minimized over prolonged periods of time. These costs might depend on the environment and diapause I and III might be better options for *Austrolebias*.

Our results demonstrate a scenario where all rates gradually become zero, and a distribution over stages is obtained that remains stable for some time. The rates of events before that point determine how many individuals there will be in each category when the cohorts are in pseudo-equilibrum. We did not find evidence of sytematically widening developmental variation leading to two alternative slow (diapause II) and fast strategies (no diapause II). However, it has not been clear which criteria were followed to assign embryos to diapause or direct-developing pathways (Furness 2015a&b) and there has been some circularity in the data analysis, as developmental process outcomes were used as explanatory variable of the same process. Data on *Austrolebias nigripinnis* in Furness et al. (2015b) suggest that developmental trajectories are not very dichotomous between embryos with direct development to a pre-hatching stage or diapause II, whereas direct developers in *Nothobranchius* have significantly larger heads). This suggests that next to presence of developmental delays, morphological embryonal life history traits have evolved differently in the different annual groups.

### Parental effects

We found a very limited number of significant effects of parental age on the survival and development of embryos. However, we did find variation between parental pairs for survival and developmental rates, which could encompass parental effects. Austrolebias annual fish do have substantial egg size variability (Moshgani and Van Dooren 2011), even when average egg size does not depend on parental age. This might be a different manner in which parental effects on survival and development might operate.

### Ptifalls

Our incubation protocol differed from incubation methods used by others, i.e., with eggs placed either on top of or under a small layer of peat moss. Alternative setups to study desiccation effects might be relevant for annual fish, but care should be taken in controlling environmental variability. In the field, it is expected that changes in soil dryness will make availability of oxygen and dryness covary, such that an experiment designed to separate oxygen and desiccation effects would be crucial to explore plasticity and survival effects of both. We incubated embryos in normoxia and with different levels of desiccation. With crossed desiccation and oxygen levels embryos might show different desiccation sensitivities depending on oxygen availability. Such a more involved experimental design would also allow a direct comparison of annuals and non-annuals, as the environmental design would include environments characteristic for either group, especially when a range of durations of desiccation are imposed. However, the amount of individuals required for such an experiment seems forbidding without automated observation of survival and developmental state.

Our data loggers did not allow us to demonstrate that we reached levels of desiccation precisely as intended, and there might be other side effects of salts than by just controlling humidity levels. The desiccators with different salts clearly smelled differently and it can be expected that molecules enter the medium with embryos as well. If such effects were there, they seem to have been limited.

We reserved the term diapause for cohorts with an absence of events for a prolonged period, as we did not use accessory individual data on their metabolism or the advance of morphological structures which could have provided measures of diapause depth at the individual level. We did record the changes in coiling direction of individual embryos finding that it occurred at a lower rate than in non-annuals. We should try to correlate different invasive and non-invasive manners of measuring metabolic activity, to see which non-invasive measures can be used as fast and reliable proxies across a wide range of killifish species in the future.

### Outlook

Timing is important in evolution, to match life histories to environments and to accelerate or delay the pace of life when advantageous. For developmental processes, this has led to the definition and study of heterochrony, an evolutionary change in the rate or timing of developmental processes (Horder 2013), or heterokairy, plasticity for developmental timing (Spicer & Burggren, 2003). Our analysis consisted of a detailed dissection of the developmental life history of embryos, and uses time-to-event analysis for analysis. The demography was followed in full, by estimating survival and rates of passing into next developemental stage. Such a dissection is important to understand life history evolution, but a lack of statistical power can occur, because not all cohorts occur in large sample sizes and the events occurring within them in certain environments cannot be controlled easily. Contrary to Furness et al (2015a&b) and Polacik et al. (2017) we did not assign individuals to alternative developmental trajectories to then redo an analysis of events that occurred before the individuals could be classified. Models such as the ones selected here could be used to estimate sensitivities of demographic parameters to environmental and genetic modification, which would allow exploring where the strongest selection gradients on developmental processes are situated.

## Conclusions

Mild desiccation and rewetting affect survival, rates of development and hatching probability in *Austrolebias bellottii*, but not the fractions of embryos that arrest development in particular stages. We can conclude that the incidences of diapause has become relatively independent of the occurrence of mild desiccation, as if they have become assimilated. In contrast to the responses observed in the non-annual rivulids, *Austrolebias* accelerates development into the pre-hatching stage in response to mild desiccation.

## Declarations

Animal Care and handling protocols were supervised by Animal Welfare committee of Leiden Faculty of Sciences and Medicine. Our data and example scripts will be made available for analysis on dryad. There are no competing interests and no funding to declare.

## Authors’ contributions

TJMVD conceived the study. IVL and TJMVD performed the experiments, analyzed the data, and wrote the manuscript. Both authors read and approved the final manuscript.

## Acknowledgements

We thank Menno Schilthuizen for encouragement, Leiden University for waiting with demolishing our aquarium facilities until we were done. Martin, Fidel and Azul Fourcade and Maria Tomjanovich we thank for hospitality while collecting *A. bellottii* lines in Ingeniero Maschwitz, Argentina.

## Supplementary Material

**Table S1.** Survival analysis per stage censored before the long interval between observations. “NS” indicates non-significant effects. Period × treatment interactions were never significant and therefore not listed. Significant effects are indicated by the Chi-squared statistic and p-value. For the salts used, those which had significantly different parameter estimates from desiccators with water are given differently shaded cells. Period effects could not be fitted in most models.

| Stage | Parent | Date | Plate | KCL | KNO3 | NH4H2PO4 | Salts | Temp | Age | Preceding |
| --- | --- | --- | --- | --- | --- | --- | --- | --- | --- | --- |
| 1 | 54.47<br>< 0.001 | 70.68<br>< 0.001 | 10.10<br>0.001 |  |  |  | NS | NS | NS |  |
| 2 | NS | 5.08<br>0.024 | NS |  |  |  | NS | NS | NS | 8.51<br>0.004 |
| 3 | NS | NS | NS |  |  |  | NS | NS | NS | NS |
| 4 | NS | NS | NS |  |  |  | NS | NS | NS | NS |
| 5 | NS | NS | NS |  |  |  | NS | NS | NS | NS |

**Table S2.**
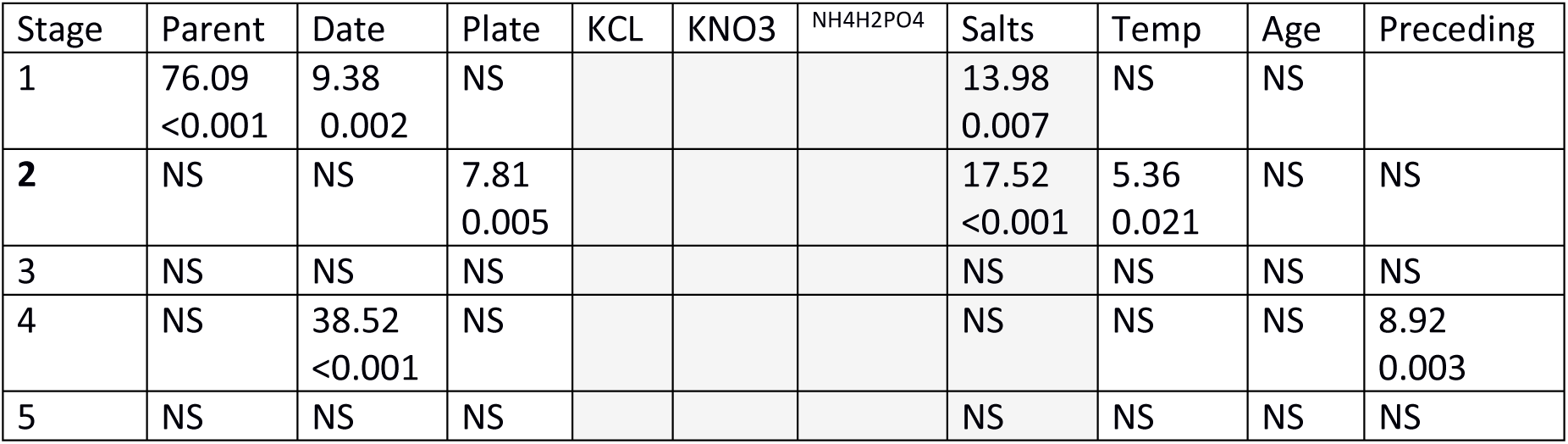
Analysis of developmental rates per stage, censored before the long interval. “NS” indicates non-significant effects. Period × treatment interactions were never significant and therefore not listed. Significant effects are indicated by Chi-squared statistic and p-value. For the salts used, those which had significantly different parameter estimates from desiccators with water are given differently shaded cells. For stages written in bold, we did not include time intervals in the rewetted period in the analysis.

| Stage | Parent | Date | Plate | KCL | KNO3 | NH4H2PO4 | Salts | Temp | Age | Preceding |
| --- | --- | --- | --- | --- | --- | --- | --- | --- | --- | --- |
| 1 | 76.09<br><0.001 | 9.38<br>0.002 | NS |  |  |  | 13.98<br>0.007 | NS | NS |  |
| **2** | NS | NS | 7.81<br>0.005 |  |  |  | 17.52<br><0.001 | 5.36<br>0.021 | NS | NS |
| 3 | NS | NS | NS |  |  |  | NS | NS | NS | NS |
| 4 | NS | 38.52<br><0.001 | NS |  |  |  | NS | NS | NS | 8.92<br>0.003 |
| 5 | NS | NS | NS |  |  |  | NS | NS | NS | NS |

**Table S3.**
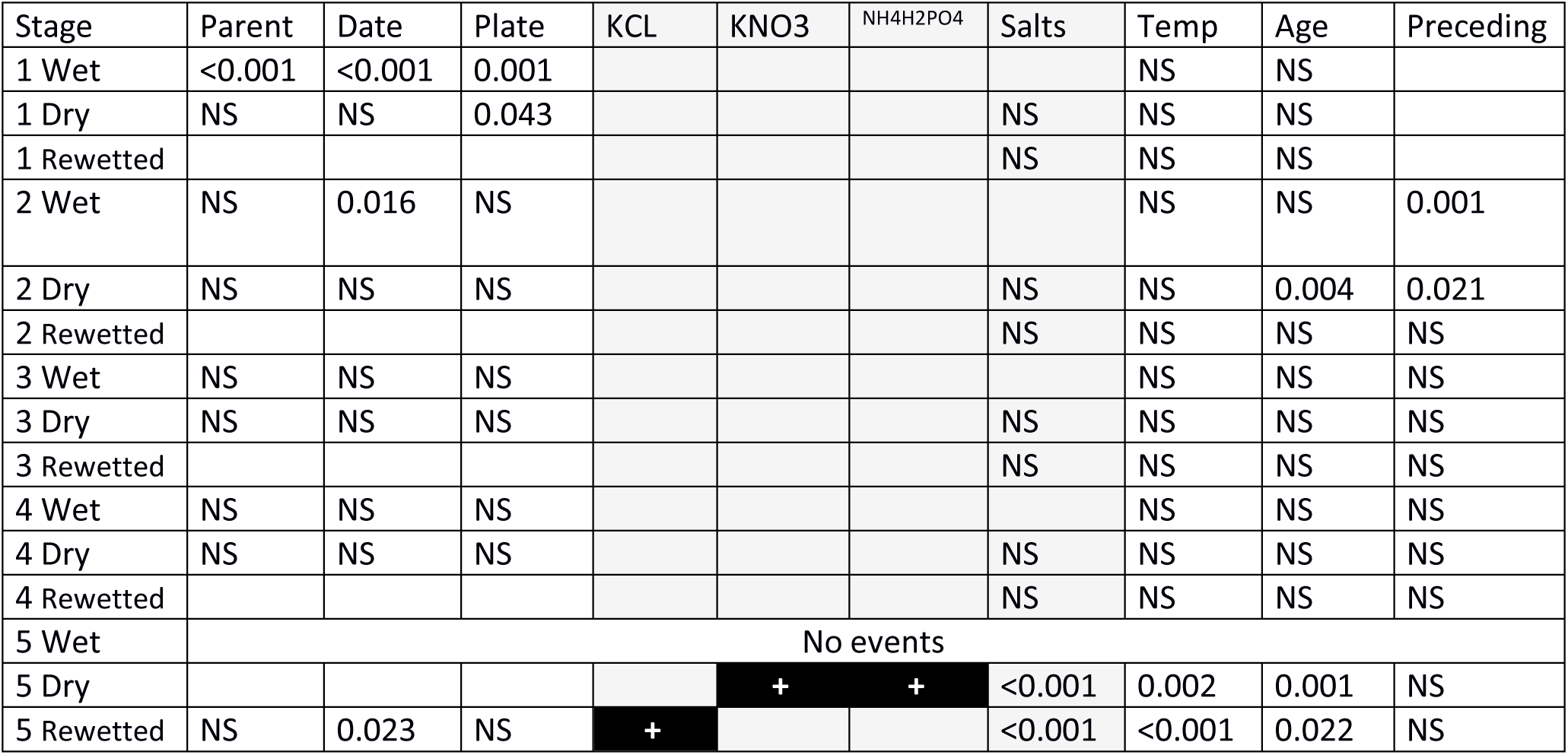
Survival analysis, per stage and period separately.. Significant effects are listed, based on likelihood ratio Chisq tests. Significant effects are indicated by p-values. When the effect was not tested, cells are left empty. For the salts used, those which had significantly different parameter estimates are given differently shaded cells.

| Stage | Parent | Date | Plate | KCL | KNO3 | NH4H2PO4 | Salts | Temp | Age | Preceding |
| --- | --- | --- | --- | --- | --- | --- | --- | --- | --- | --- |
| 1 Wet | <0.001 | <0.001 | 0.001 |  |  |  |  | NS | NS |  |
| 1 Dry | NS | NS | 0.043 |  |  |  | NS | NS | NS |  |
| 1 Rewetted |  |  |  |  |  |  | NS | NS | NS |  |
| 2 Wet | NS | 0.016 | NS |  |  |  |  | NS | NS | 0.001 |
| 2 Dry | NS | NS | NS |  |  |  | NS | NS | 0.004 | 0.021 |
| 2 Rewetted |  |  |  |  |  |  | NS | NS | NS | NS |
| 3 Wet | NS | NS | NS |  |  |  |  | NS | NS | NS |
| 3 Dry | NS | NS | NS |  |  |  | NS | NS | NS | NS |
| 3 Rewetted |  |  |  |  |  |  | NS | NS | NS | NS |
| 4 Wet | NS | NS | NS |  |  |  |  | NS | NS | NS |
| 4 Dry | NS | NS | NS |  |  |  | NS | NS | NS | NS |
| 4 Rewetted |  |  |  |  |  |  | NS | NS | NS | NS |
| 5 Wet | No events |  |  |  |  |  |  |  |  |  |
| 5 Dry |  |  |  |  | + | + | <0.001 | 0.002 | 0.001 | NS |
| 5 Rewetted | NS | 0.023 | NS | + |  |  | <0.001 | <0.001 | 0.022 | NS |

**Table S4.** Development per period and per stage. Significant effects are indicated by *p*-values. When the effect was not tested, cells are left empty. For the salts used, those which had significantly different parameter estimates are given differently shaded cells.

| Stage | Parent | Date | Plate | KCL | KNO3 | NH4H2PO4 | Salts | Temp | Age | Preceding |
| --- | --- | --- | --- | --- | --- | --- | --- | --- | --- | --- |
| 1 Wet | <0.001 | 0.006 | NS |  |  |  |  | NS | NS |  |
| 1 Dry | 0.004 | 0.001 | 0.027 |  |  |  | NS | NS | NS |  |
| 1 Rewetted |  |  |  |  |  |  | NS | NS | NS |  |
| 2 Wet | NS | 0.001 | 0.033 |  |  |  |  | NS | NS | 0.003 |
| 2 Dry | NS | NS | NS |  |  |  | NS | NS | NS | 0.001 |
| 2 Rewetted |  |  |  |  |  |  | NS | NS | NS | NS |
| 3 Wet | NS | NS | NS |  |  |  |  | NS | NS | NS |
| 3 Dry | NS | NS | 0.036 |  |  |  | NS | NS | NS | NS |
| 3 Rewetted |  |  |  |  |  |  | NS | NS | NS | NS |
| 4 Wet | NS | 0.032 | NS |  |  |  |  | 0.027 | NS | <0.001 |
| 4 Dry | 0.016 | <0.001 | NS |  |  |  | 0.010 | NS | NS | 0.024 |
| 4 Rewetted |  |  |  |  |  |  | NS | NS | NS | NS |
| 5 Wet | NS | NS | NS |  |  |  |  | NS | NS | NS |
| 5 Dry | NS | NS | NS |  |  |  | NS | NS | NS | NS |
| 5 Rewetted | No events |  |  |  |  |  |  |  |  |  |

**Fig Surv1.**
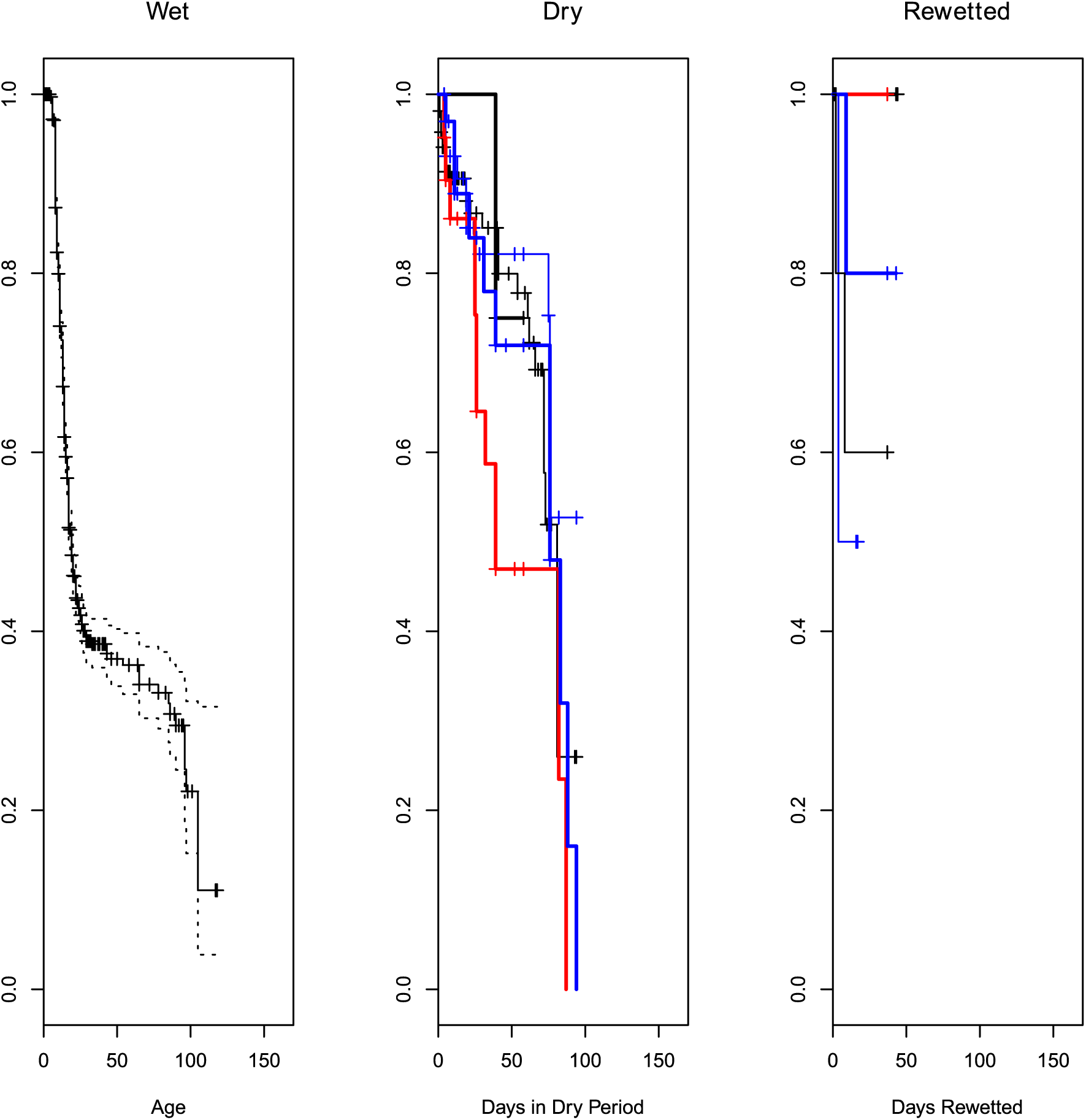
Survival stage one different periods. From left to right: before desiccation, during desiccation, and when water was added again. In the two rightmost panels age is rescaled to the time where the regime started (i.e. start of dry period, rewetted). The control group is added to each panel as well, with the average age at which the regime started substracted from the actual age of embryos. This was done to facilitate comparisons between treatments. Black, thin: Control group, blue, thin: dessicator with H20, red thick KNO3 blue thick NH4H2PO4 black, thick: KCL.

**Fig Surv2.**
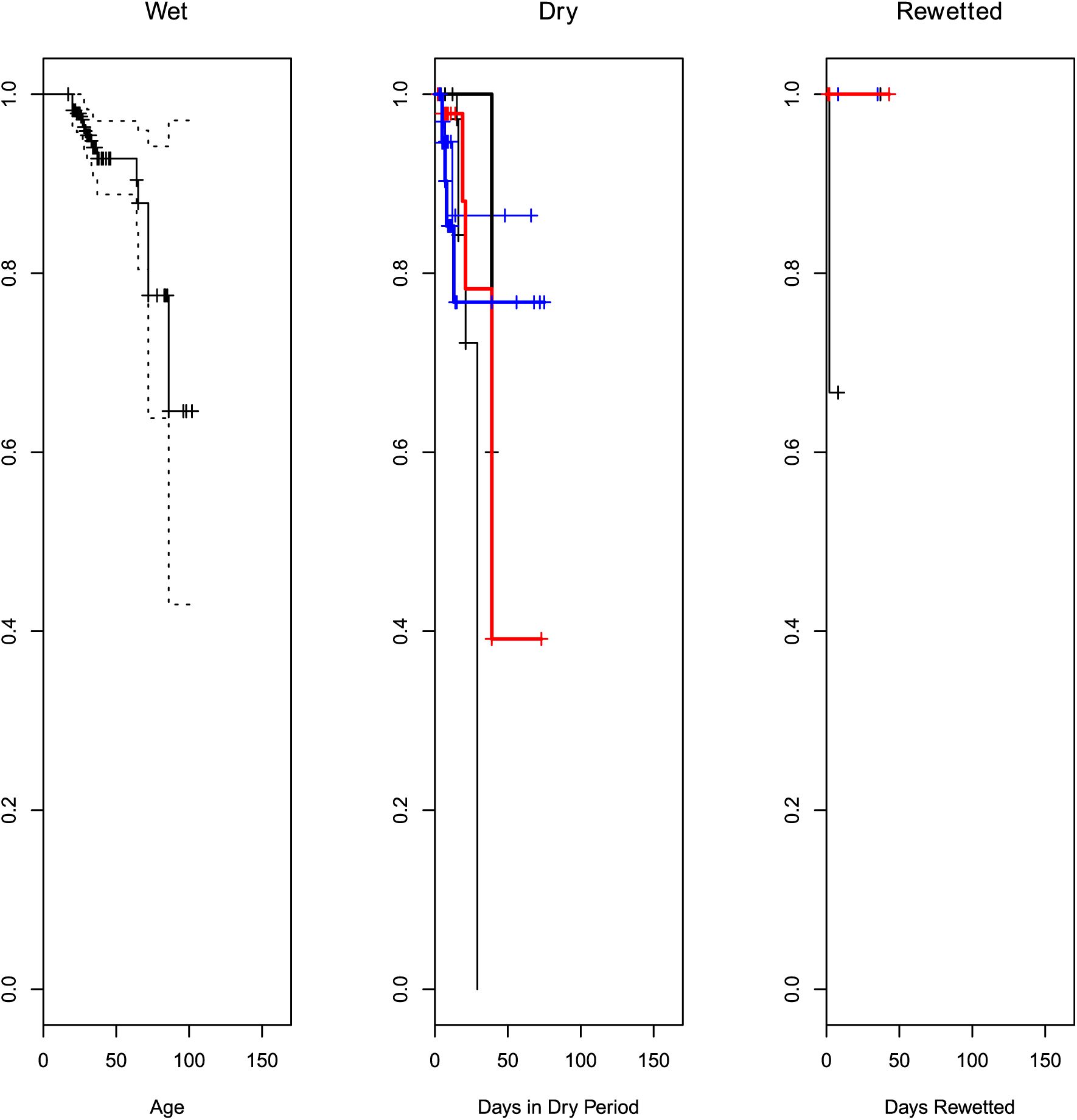
Survival stage two different periods. From left to right: before desiccation, during desiccation, and when water was added again. In the two rightmost panels age is rescaled to the time where the regime started (i.e. start of dry period, rewetted). The control group is added to each panel as well, with the average age at which the regime started substracted from the actual age of embryos. This was done to facilitate comparisons between treatments. Black, thin: Control group, blue, thin: dessicator with H20, red thick KNO3 blue thick NH4H2PO4 black, thick: KCL.

**Fig Surv3.**
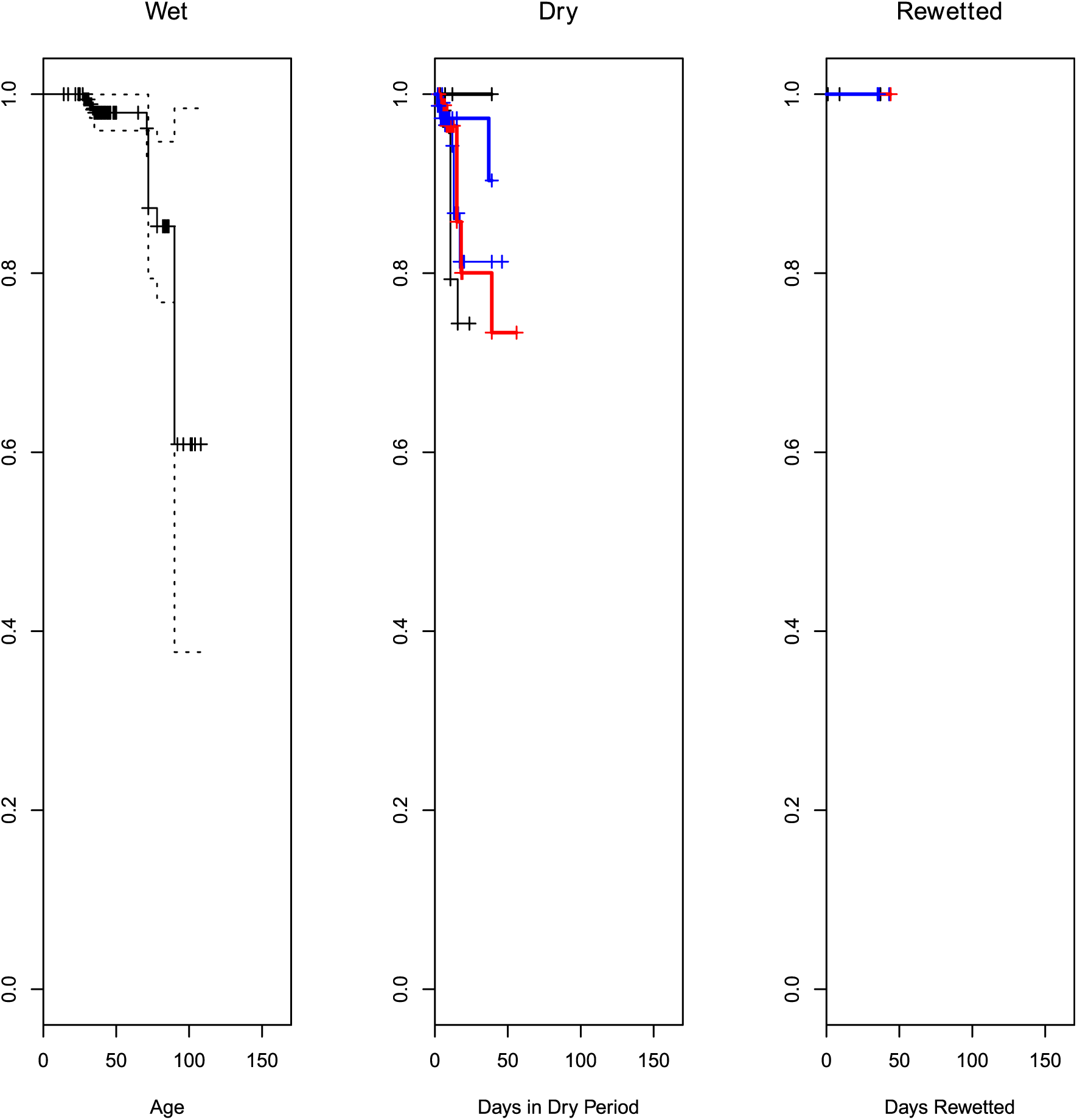
Survival stage three in different periods. From left to right: before desiccation, during desiccation, and when water was added again. In the two rightmost panels age is rescaled to the time where the regime started (i.e. start of dry period, rewetted). The control group is added to each panel as well, with the average age at which the regime started substracted from the actual age of embryos. This was done to facilitate comparisons between treatments. Black, thin: Control group, blue, thin: dessicator with H20, red thick KNO3 blue thick NH4H2PO4 black, thick: KCL.

**Fig Surv4.**
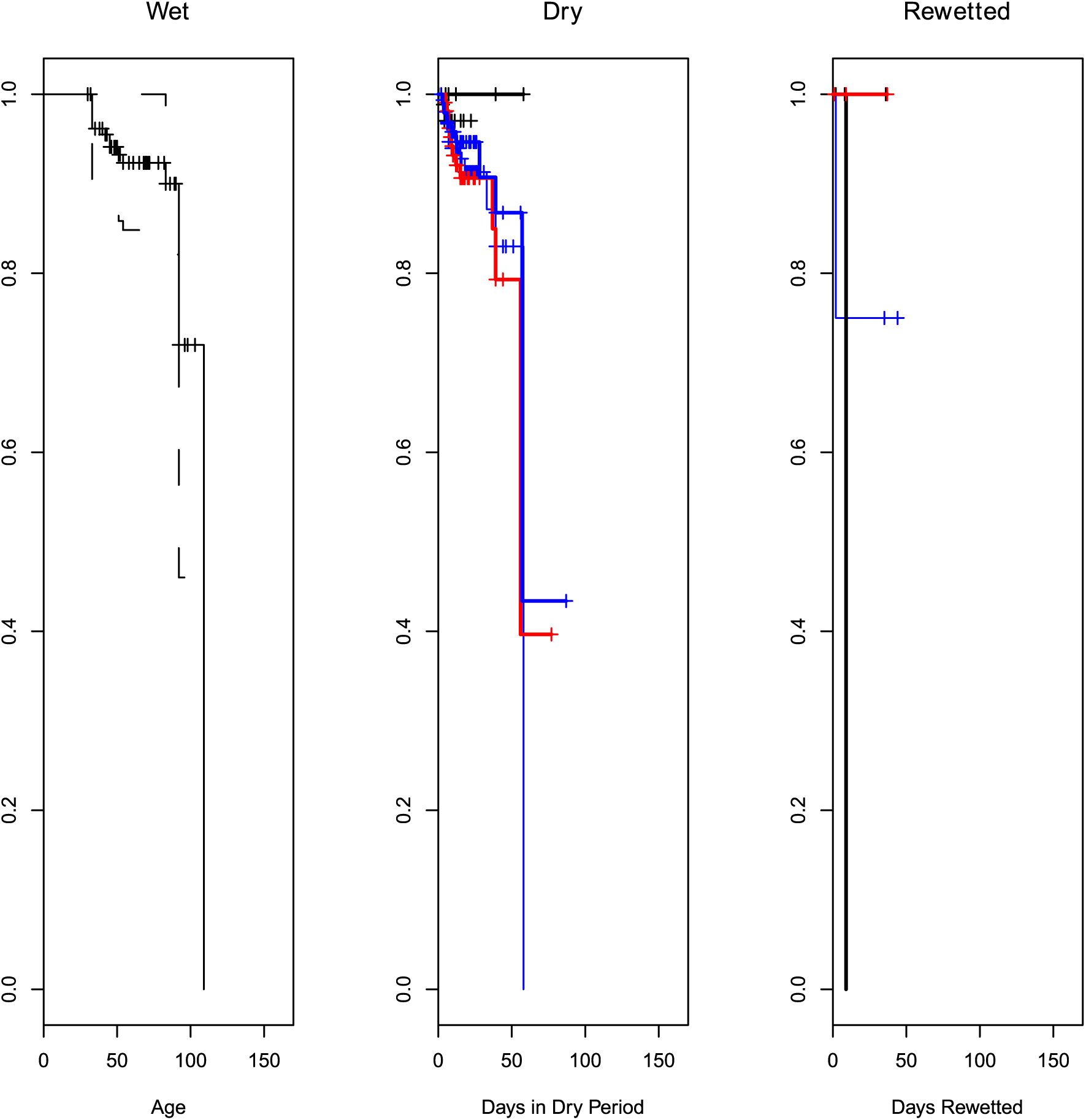
Survival stage four different periods. From left to right: before desiccation, during desiccation, and when water was added again. In the two rightmost panels age is rescaled to the time where the regime started (i.e. start of dry period, rewetted). The control group is added to each panel as well, with the average age at which the regime started substracted from the actual age of embryos. This was done to facilitate comparisons between treatments. Black, thin: Control group, blue, thin: dessicator with H20, red thick KNO3 blue thick NH4H2PO4 black, thick: KCL.

**Fig Surv5.**
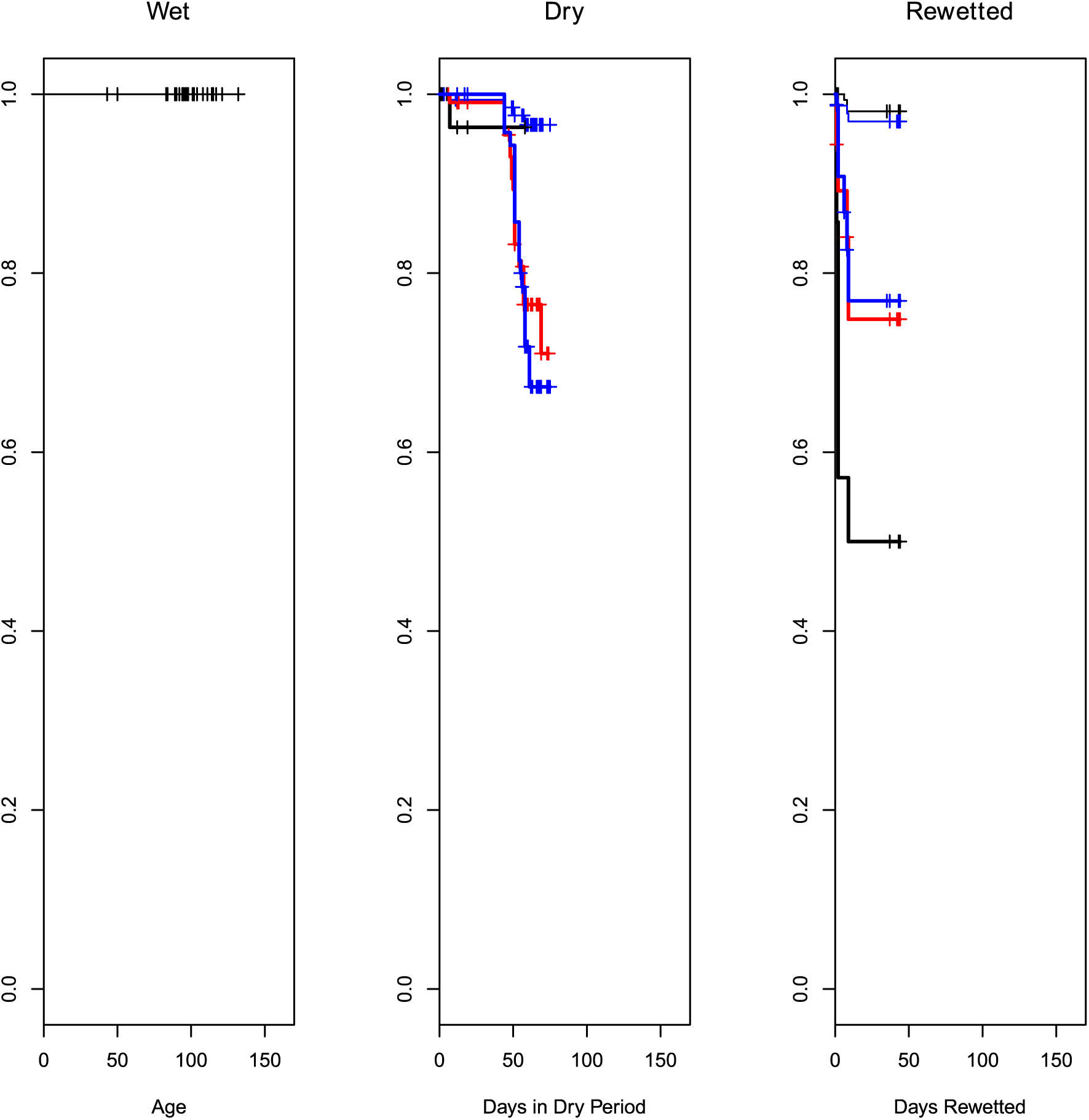
Survival stage five in different periods. From left to right: before desiccation, during desiccation, and when water was added again. In the two rightmost panels age is rescaled to the time where the regime started (i.e. start of dry period, rewetted). The control group is added to each panel as well, with the average age at which the regime started substracted from the actual age of embryos. This was done to facilitate comparisons between treatments. Black, thin: Control group, blue, thin: dessicator with H20, red thick KNO3 blue thick NH4H2PO4 black, thick: KCL.

**Fig Dev1.**
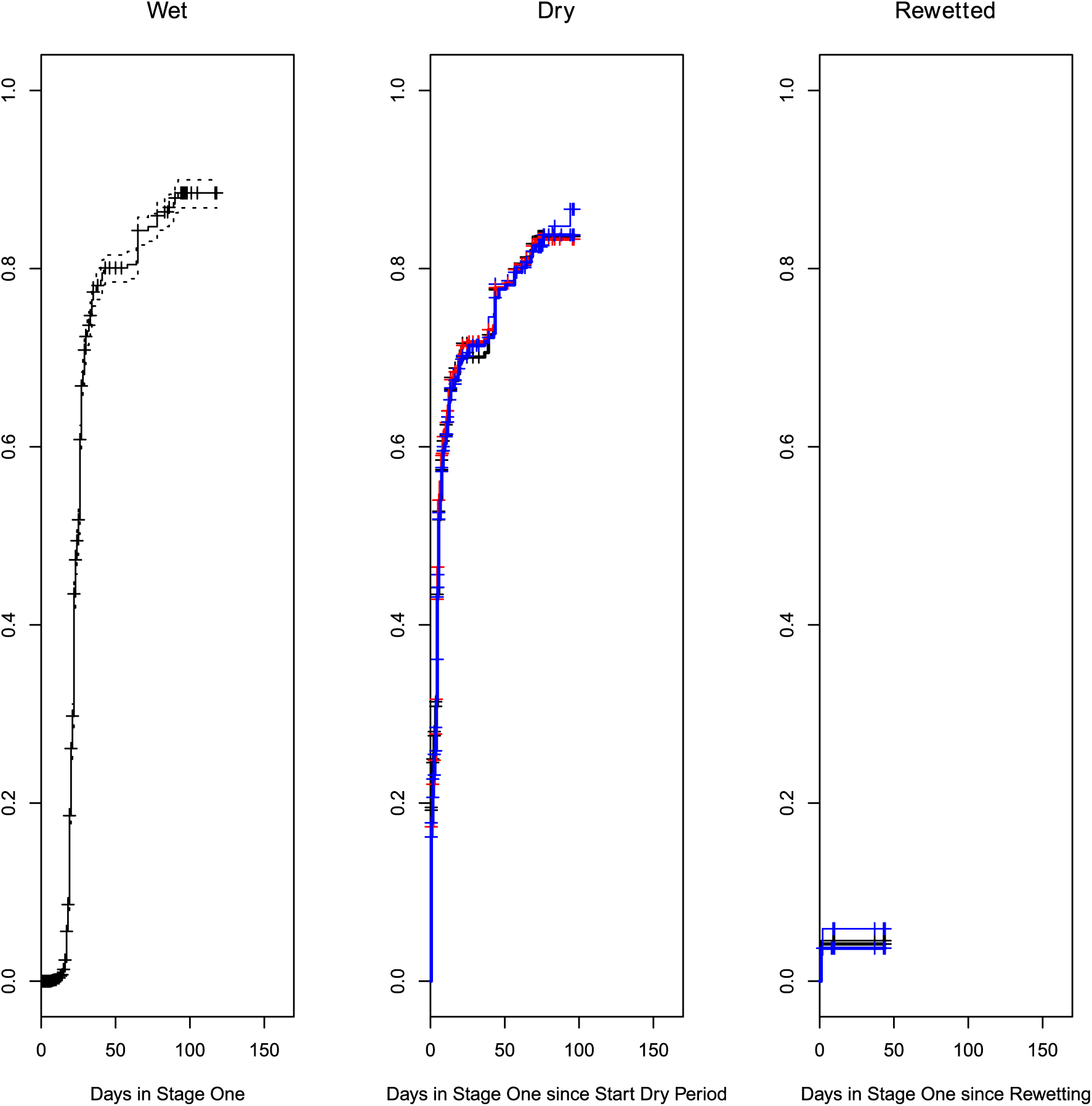
Development from stage one into stage two in the different periods in the experiment. On the y-axis is plotted the fraction in the population that made the transition. From left to right: before desiccation, during desiccation, and when water was added again. In the two rightmost panels age-within-stage is rescaled to the time where the regime started (i.e. start of dry period, rewetted). The control group is added to each panel as well, with the average age at which the regime started substracted from the actual age-within-stage of embryos. This was done to facilitate comparisons between treatments. Black, thin: Control group, blue, thin: dessicator with H20, red thick KNO3 blue thick NH4H2PO4 black, thick: KCL.

**Fig Dev2.**
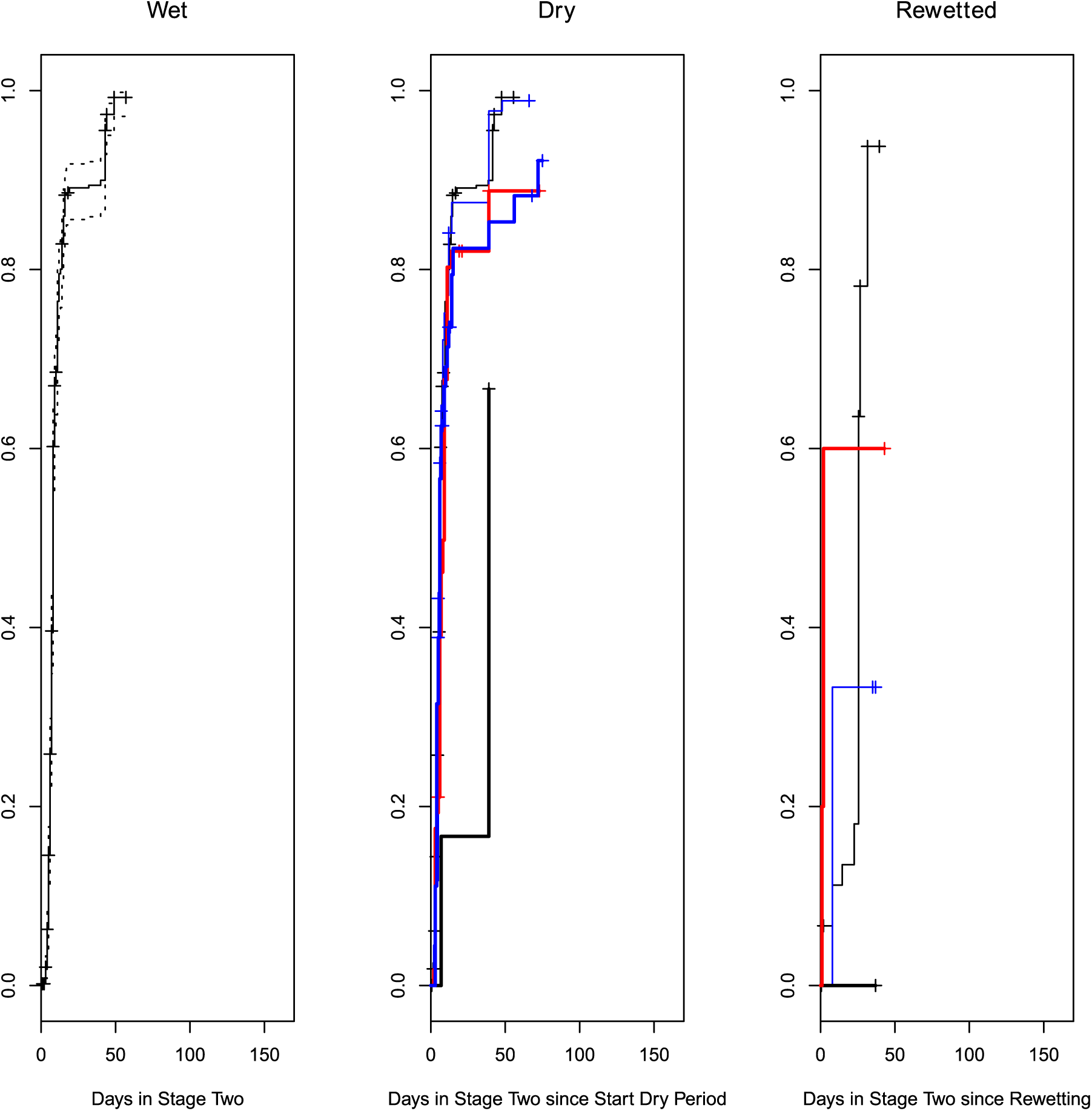
Development from stage two into stage three in the different periods in the experiment. From left to right: before desiccation, during desiccation, and when water was added again. In the two rightmost panels age-within-stage is rescaled to the time where the regime started (i.e. start of dry period, rewetted). The control group is added to each panel as well, with the average age at which the regime started substracted from the actual age-within-stage of embryos. This was done to facilitate comparisons between treatments. Black, thin: Control group, blue, thin: dessicator with H20, red thick KNO3 blue thick NH4H2PO4 black, thick: KCL.

**Fig Dev3.**
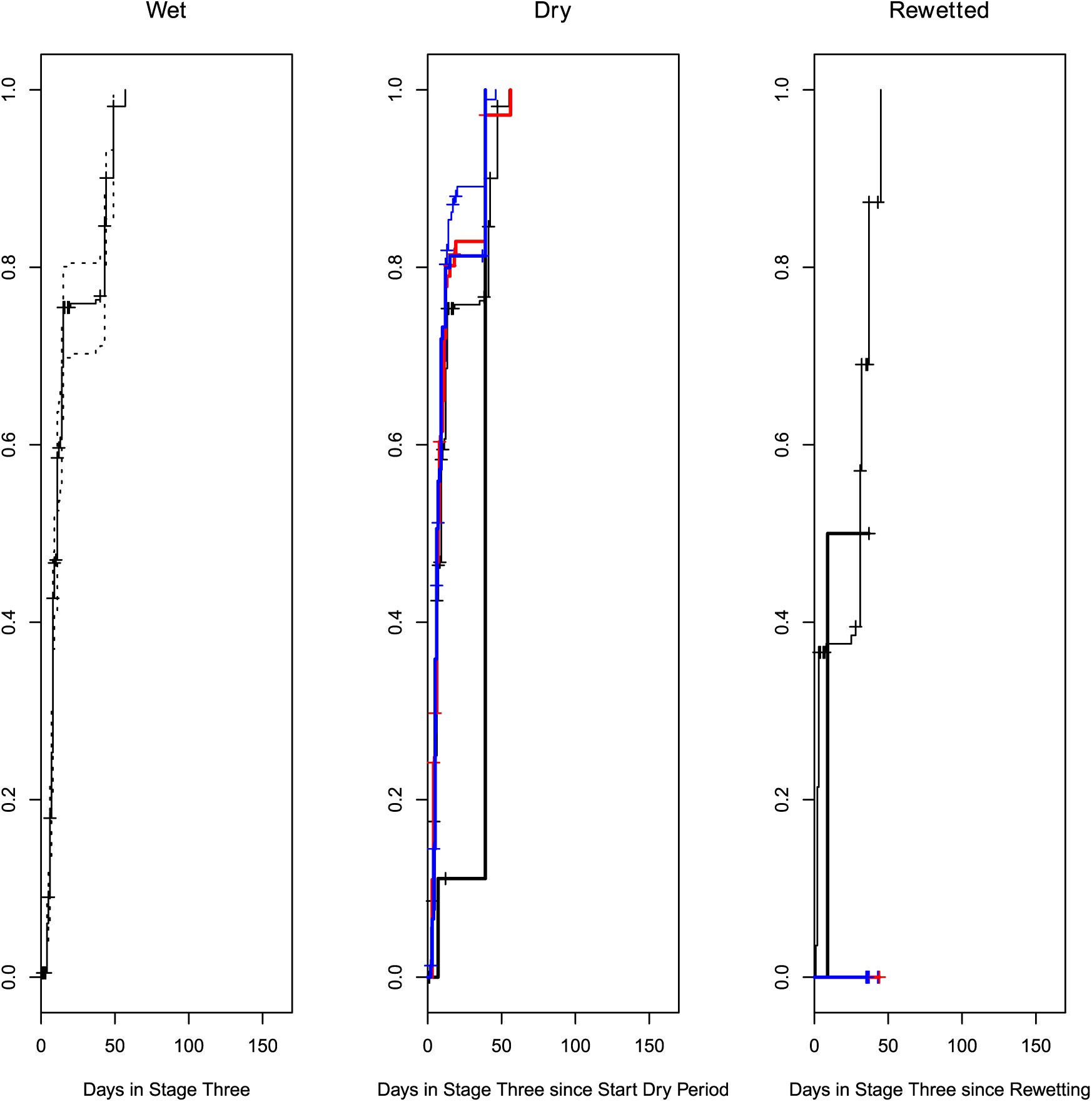
Development from stage three into stage four in the different periods in the experiment. From left to right: before desiccation, during desiccation, and when water was added again. In the two rightmost panels age-within-stage is rescaled to the time where the regime started (i.e. start of dry period, rewetted). The control group is added to each panel as well, with the average age at which the regime started substracted from the actual age-within-stage of embryos. This was done to facilitate comparisons between treatments. Black, thin: Control group, blue, thin: dessicator with H20, red thick KNO3 blue thick NH4H2PO4 black, thick: KCL.

**Fig Dev4.**
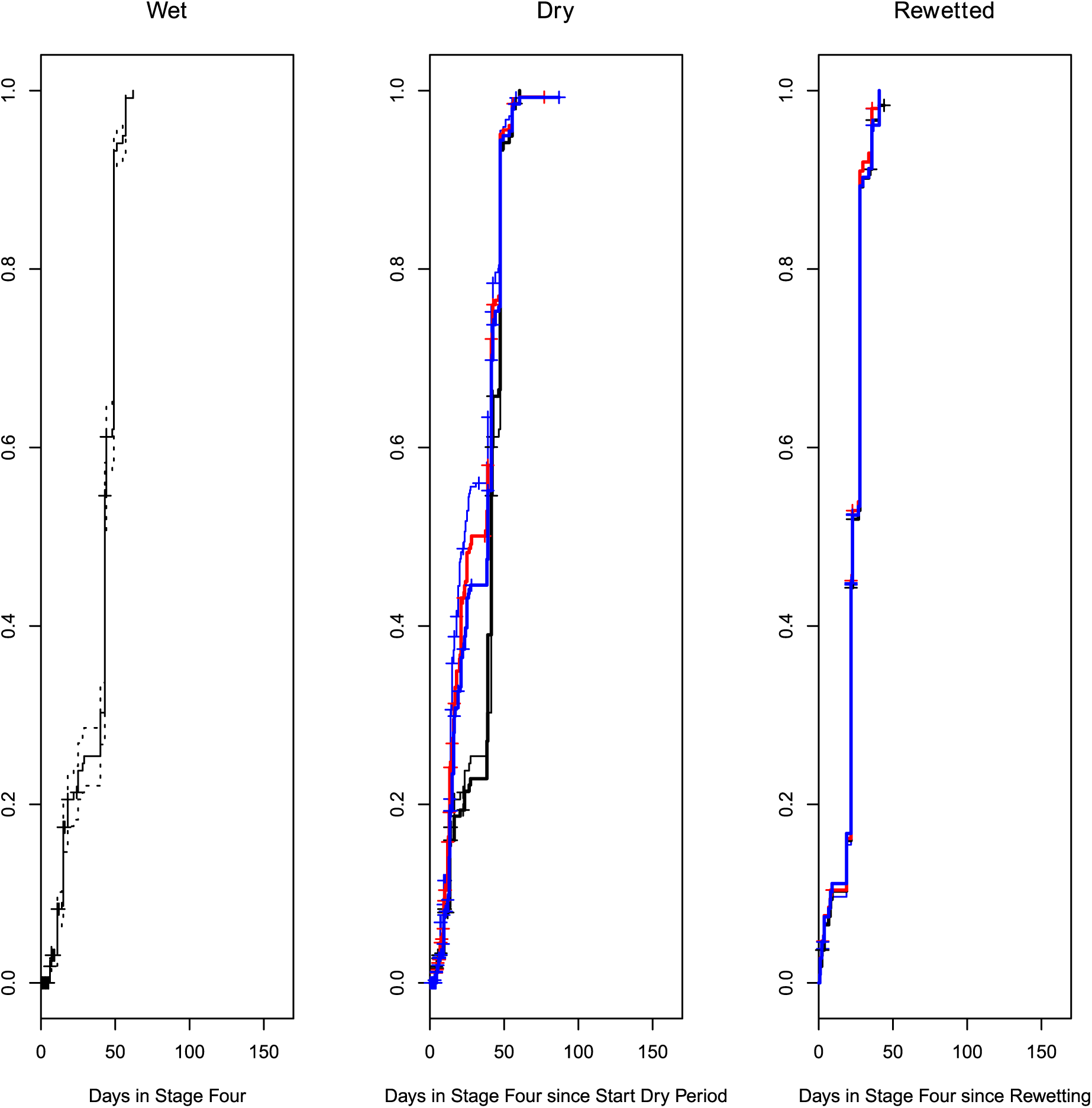
Development from stage four into stage five in the different periods in the experiment. From left to right: before desiccation, during desiccation, and when water was added again. In the two rightmost panels age-within-stage is rescaled to the time where the regime started (i.e. start of dry period, rewetted). The control group is added to each panel as well, with the average age at which the regime started substracted from the actual age-within-stage of embryos. This was done to facilitate comparisons between treatments. Black, thin: Control group, blue, thin: dessicator with H20, red thick KNO3 blue thick NH4H2PO4 black, thick: KCL.

**Fig Dev5.**
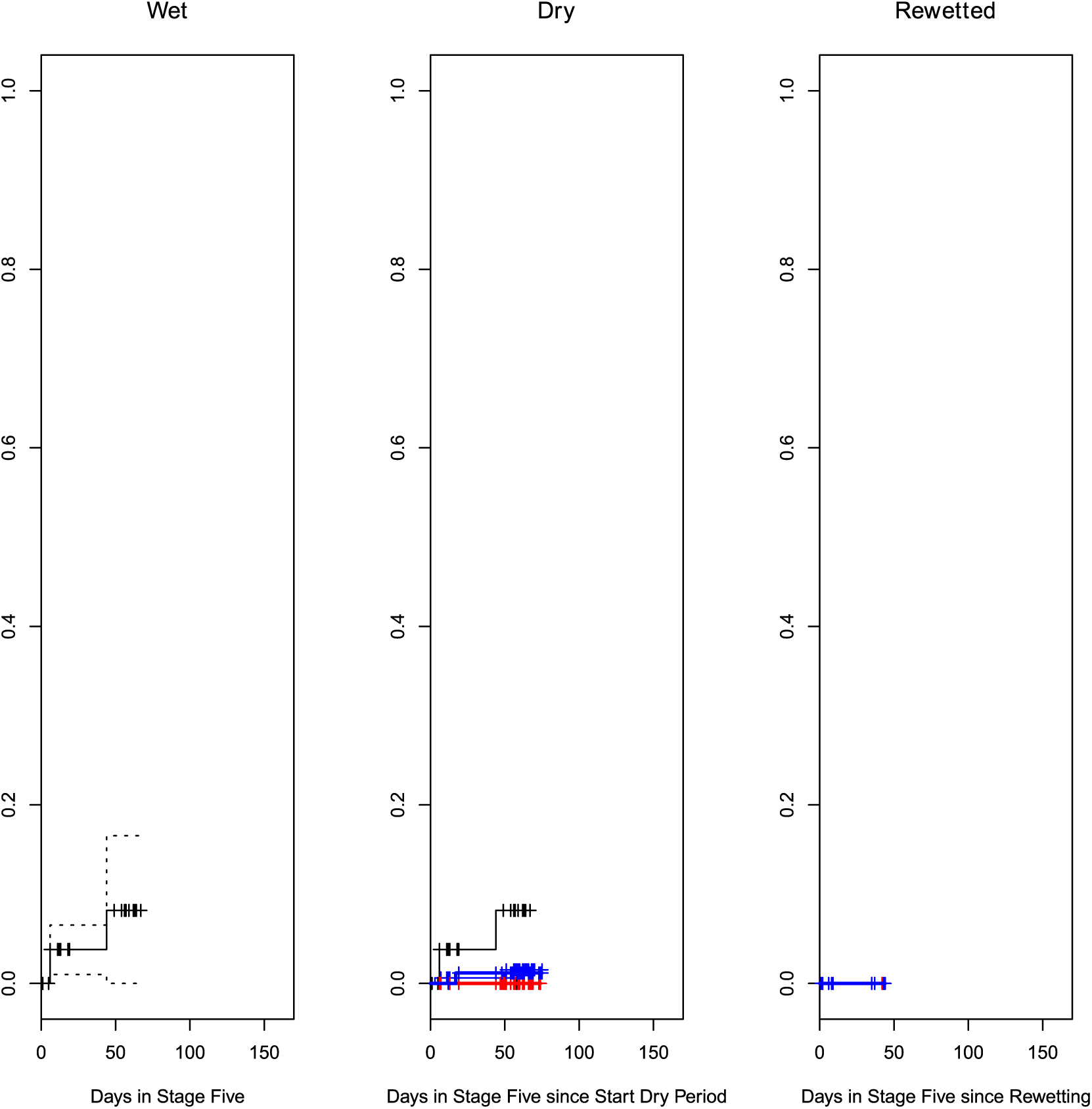
Proportions of individuals in stage five that do not hatch spontaneously. From left to right: before desiccation, during desiccation, and when water was added again. In the two rightmost panels age-within-stage is rescaled to the time where the regime started (i.e. start of dry period, rewetted). The control group is added to each panel as well, with the average age at which the regime started substracted from the actual age-within-stage of embryos. This was done to facilitate comparisons between treatments. Black, thin: Control group, blue, thin: dessicator with H20, red thick KNO3 blue thick NH4H2PO4 black, thick: KCL.

